# Long-term immune reprogramming of classical monocytes with altered ontogeny mediates enhanced lung injury in sepsis survivors

**DOI:** 10.1101/2025.05.16.654442

**Authors:** Scott J. Denstaedt, Breanna McBean, Alan P Boyle, Brett C. Arenberg, Matthias Mack, Bethany B. Moore, Michael W. Newstead, Yanmei Deng, Alexey I. Nesvizhskii, Benjamin H. Singer, Jennifer Cano, Hallie C. Prescott, Helen S. Goodridge, Rachel L. Zemans

## Abstract

Patients who survive sepsis are predisposed to new hospitalizations for respiratory failure, but the underlying mechanisms are unknown. Using a murine model in which prior sepsis predisposes to enhanced lung injury, we previously discovered that classical monocytes persist in the lungs after long-term recovery from sepsis and exhibit enhanced cytokine expression after secondary challenge with intra-nasal lipopolysaccharide. Here, we hypothesized that immune reprogramming of post-sepsis monocytes and altered ontogeny predispose to enhanced lung injury. Monocyte depletion and/or adoptive transfer was performed three weeks and three months after sepsis. Monocytes from post-sepsis mice were necessary and sufficient for enhanced LPS-induced lung injury and promoted neutrophil degranulation. Prior sepsis enhanced JAK-STAT signaling and AP-1 binding in monocytes and shifted monocytes toward the neutrophil-like monocyte lineage. In human sepsis and/or pneumonia survivors, monocytes were predictive of 90-day mortality and exhibit transcriptional and proteomic neutrophil-like signatures. We conclude that sepsis reprograms monocytes into a pro-inflammatory phenotype and skews bone marrow progenitors and monocytes toward the neutrophil-like lineage, predisposing to neutrophil degranulation and lung injury.

## Introduction

Nearly 40 million people survive sepsis each year^1^, yet half of these survivors are rehospitalized or die within the following year^2–6^. One in 20 sepsis survivors are rehospitalized with new lung injury, including conditions such as aspiration pneumonitis and exacerbation of chronic respiratory disease^2,4^. Our inability to develop effective strategies to prevent these complications stems from a limited understanding of the underlying mechanisms driving post-sepsis lung injury. Elevated inflammatory markers, such as circulating interleukin-6 (IL-6), C-reactive protein (CRP), and leukocyte counts, at hospital discharge, have been associated with an increased risk of rehospitalization and mortality^7–10^, suggesting that persistent immune dysregulation after sepsis may play a critical role^11^.

To elucidate the mechanisms of lung injury in sepsis survivors, we previously developed model of sterile lung injury in mice that have survived sepsis. In this model, cecal ligation and puncture (CLP) predisposes to enhanced lung injury in response to intranasal (i.n.) lipopolysaccharide (LPS) administered 3 weeks after CLP^12^. We discovered that monocytosis persists in the lungs for at least 3 weeks following sepsis alone, and these Ly6C^hi^ monocytes produced higher levels of *Tnf* in response to i.n. LPS than monocytes in mice without prior sepsis^12^. However, whether monocytes play a causal role in the enhanced lung injury induced by prior sepsis and the mechanisms through which sepsis might reprogram monocytes to exacerbate lung injury has not been studied.

Monocytes are known to play a direct role in regulating acute lung injury in the absence of prior sepsis. Ly6C^hi^ monocytes can either promote or mitigate lung injury depending on context^13–20^. For instance, in *Klebsiella pneumoniae* infection, monocytes recruited to the lung are protective, whereas in influenza infection they are detrimental^14–19,21^. These observations suggest that monocytes assume diverse functional states and have led to a paradigm whereby the inflammatory milieu of the lung during injury dictates the activation status and function of recruited monocytes. This prevailing view relies on the assumption that circulating monocytes are a homogeneous population and also fails to consider how prior exposures may modulate recruitable monocyte functions or fates.

It is well established that infection can elicit durable reprogramming of immune cells and their progenitors in the bone marrow (BM), leading to enhanced (i.e., primed or trained) or suppressed (i.e., tolerant) responses to secondary stimuli^22–26^. This reprogramming occurs via changes in transcriptional and epigenetic states, which may persist for weeks to months^24,27–32^. Such reprogramming typically involves activation or suppression of proinflammatory pathways. However, the ways in which cells are reprogrammed and the contexts in which immune reprogramming results in beneficial (e.g., host defense) or detrimental (e.g., tissue injury) responses to subsequent insults are vastly underexplored. Specifically, the extent to which respiratory complications in sepsis survivors are attributable to immune reprogramming, the specific cells that are reprogrammed, the mechanisms through which reprogrammed cells predispose to enhanced lung injury, and whether reprogramming may extend beyond alterations in inflammatory pathways are not known.

An emerging literature has revealed that BM Ly6C^hi^ monocytes comprise a heterogenous population of transcriptionally unique subsets^33–36^, including those with distinct neutrophil-like and dendritic cell (DC)-like gene expression profiles, which are pre-determined during hematopoiesis. Critically, although single-cell RNA sequencing (scRNAseq) has characterized the transcriptomes of these monocyte subsets, aside from isolated studies^37,38^, the specific functions of these subsets in homeostasis and disease remain unexplored. Specifically, the role of these novel monocyte subsets in acute lung injury and their contribution to the observed functional heterogeneity of monocytes in this context have not been investigated. Moreover, whether immune reprogramming may involve not only the activation or suppression of proinflammatory pathways but also the reprogramming of mature monocytes and progenitors towards a particular lineage, thereby influencing responses to subsequent stimuli, has not been studied.

In this study, we address key questions regarding the functional heterogeneity and immune reprogramming of monocytes. We hypothesized that Ly6C^hi^ monocytes and/or their progenitors in the BM are reprogrammed by sepsis, thereby mediating enhanced post-sepsis lung injury. Through investigations of mouse models and human disease, we discovered that sepsis induces reprogramming of BM monocytes and their progenitors, leading to activation of proinflammatory pathways, altered hematopoiesis with a shift in progenitor cell fate towards the neutrophil-like monocyte lineage, and enhanced lung injury. Our findings introduce new fundamental insights into the functions of novel monocyte subsets and demonstrate that immune reprogramming can include not only changes in activation state but also shifts in ontogeny. Furthermore, by demonstrating that reprogramming of monocytes and their progenitors mediates the enhanced lung injury observed after sepsis, we challenge the paradigm that the lung milieu dictates functional monocyte responses during lung injury. Finally, we identify novel therapeutic targets to reduce long-term pulmonary complications resulting in rehospitalization and mortality among sepsis survivor patients.

## Results

### Depletion of Ly6C^hi^ monocytes from post-CLP mice attenuates the lung injury response to LPS

We previously demonstrated that sepsis induced by CLP in mice induces persistent lung monocytosis at three weeks (3-wk) and, in response to subsequent i.n. LPS challenge, enhanced lung injury and *Tnf* expression by monocytes^12^. However, whether monocytes play a functional role in this enhanced lung injury is unknown. To determine whether Ly6C^hi^ monocytes are required for enhanced lung injury in 3-wk post-CLP mice, we depleted monocytes using an anti-CCR2 antibody (αCCR2) administered prior to i.n. LPS (Figure 1A). Flow cytometric analysis of exudate cells in bronchoalveolar lavage (BAL) confirmed effective depletion of monocytes and monocyte-derived exudate macrophages (Figures S1, S2). Post-CLP mice treated with αCCR2 antibody (post-CLP_αCCR2_) had significantly reduced BAL albumin, indicating decreased lung permeability, at 72 hours after LPS relative to post-CLP mice treated with isotype control antibody (post-CLP_ISO_) (Figure 1B). Importantly, in contrast to the protective effect of monocyte depletion on lung injury after CLP, monocyte depletion in unoperated control mice demonstrated a trend towards exacerbation of lung injury, as previously reported^14^ (Figure 1C). Accordingly, there was less alveolar epithelial injury, as suggested by BAL RAGE^39^ levels in post-CLP_αCCR2_ relative to post-CLP_ISO_ mice (Figure 1D). Neutrophil recruitment to the airspaces was unchanged (Figure 1E). Monocyte depletion significantly reduced BAL IL-6 levels (Figure 1F) but had little effect on other inflammatory mediators (Figure S3A-C). Taken together, these data demonstrate that monocytes mediate enhanced lung injury following LPS challenge in post-CLP mice, in contrast to their protective role in lung injury in unoperated mice.

**Figure 1.**
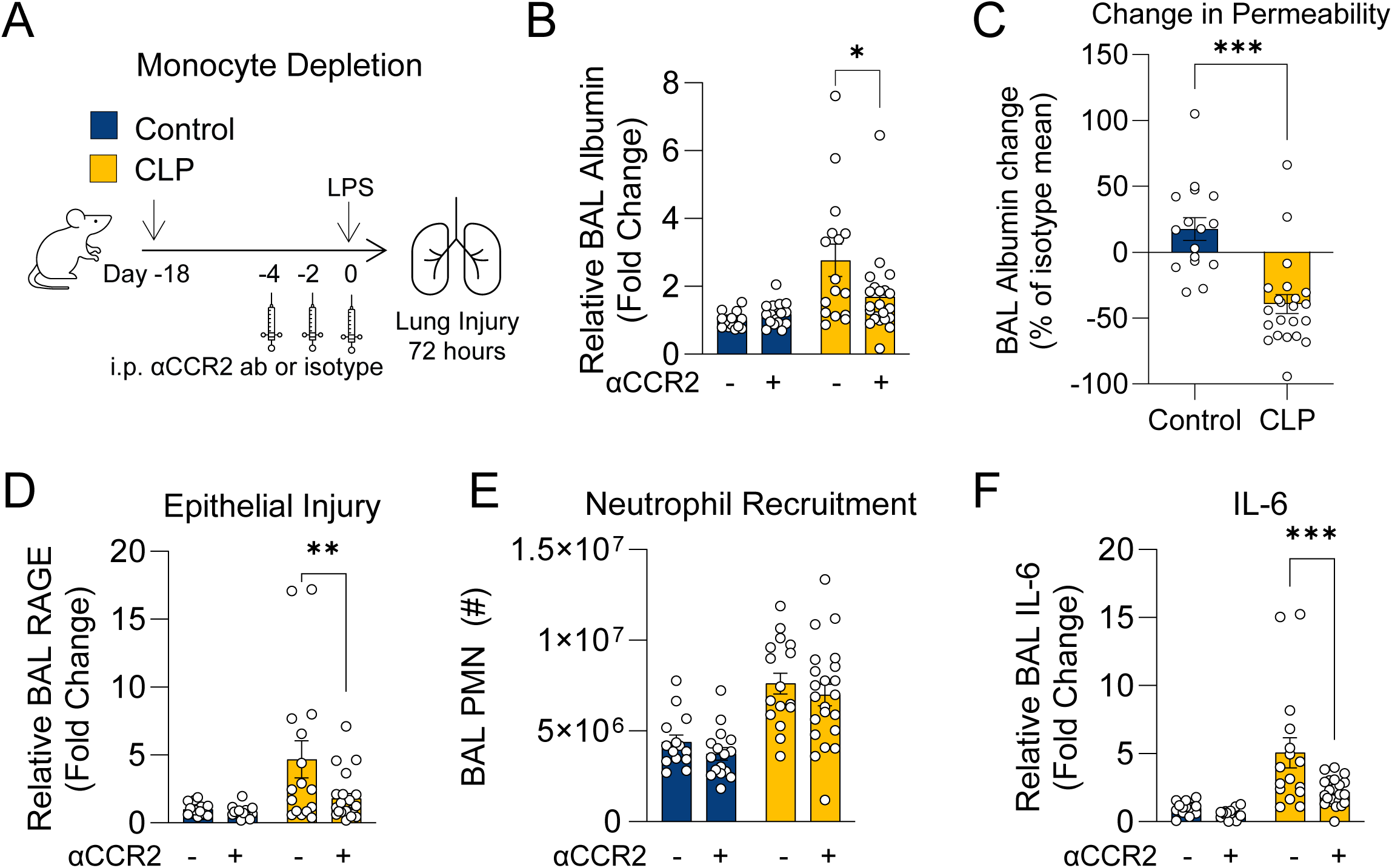
Monocyte depletion improves lung injury and inflammation in post-CLP mice. (A) Experimental design for monocyte depletion with anti-CCR2 antibody (αCCR2) or isotype prior to i.n. LPS. Lung permeability assessed by BAL albumin (B), change in permeability relative to isotype-treated for each condition (C), epithelial injury assessed by BAL RAGE (D), neutrophil recruitment (E), and BAL IL-6 (F) shown 72 hours after i.n. LPS. n = 4-8 per group, 3 cohorts; 2 of 18 mice in the iso-treated CLP group died by 72 hours, no other deaths were recorded. BAL protein levels expressed relative to the mean of isotype-treated unoperated control mice. Mean ± SEM, Sidak post-hoc p-value (B, D-F), Welch’s t-test (C) shown. * p < 0.05, ** p < 0.01, *** p < 0.001. CLP, Cecal Ligation and Puncture. LPS, Lipopolysaccharide. BAL, bronchoalveolar lavage. RAGE, receptor for advanced glycation end-products. PMN, polymorphonuclear.

### Adoptive transfer of Ly6C^hi^ monocytes from post-CLP mice fail to protect against LPS-induced lung injury

Having shown that post-CLP monocytes are necessary for enhanced lung injury after LPS, we aimed to determine whether 3-wk post-CLP monocytes are sufficient to exacerbate lung injury in response to LPS. Ly6C^hi^ monocytes from the BM of post-CLP or unoperated control mice were adoptively transferred to naïve *Ccr2^-/-^*mice, followed by i.n. LPS administration (Figure 2A). Use of *Ccr2^-/-^* mice isolates the effect of the adoptively transferred *Ccr2^wt^* monocytes, eliminating any potential confounding effects of endogenous recruited monocytes in the recipient mice. Consistent with prior literature^14^ and our monocyte depletion experiments (Figure 1), transfer of *Ccr2^wt^* monocytes harvested from unoperated control mice tended to decrease lung permeability, as measured by BAL albumin, compared to transfer of *Ccr2^-/-^*monocytes (Figure 2B). In contrast, post-CLP monocyte transfer resulted in increased BAL albumin and total protein relative to *Ccr2^wt^*unoperated control monocytes (Figure 2B,C). There were no differences in neutrophil recruitment to the airspace (Figure 2D) or in levels of most cytokines and chemokines (Figure 2E, Figure S4). Collectively, Figures 1 and 2 show that post-CLP Ly6C^hi^ monocytes are detrimental in LPS-induced lung injury in contrast to the more protective functional program of unoperated control Ly6C^hi^ monocytes, suggesting that CLP reprograms monocytes from a protective phenotype to an injurious phenotype.

**Figure 2.**
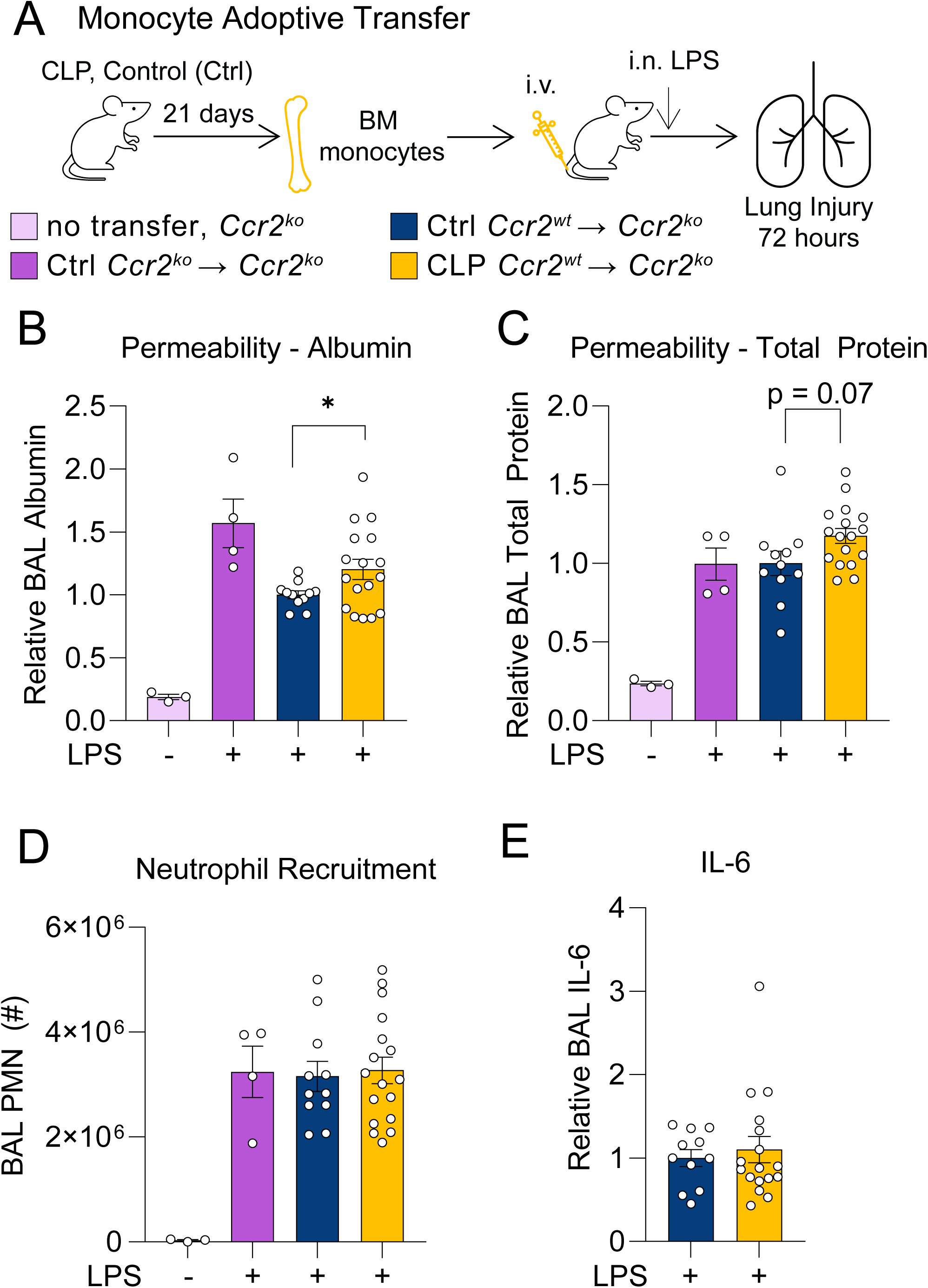
Adoptive transfer of post-CLP monocytes enhances the lung injury response to LPS. Bone marrow Ly6C^hi^ monocytes from *Ccr2^-/-^*, *Ccr2^wt^*age-matched unoperated control, and *Ccr2^wt^* 3-wk post-CLP mice were isolated and administered into *Ccr2^-/-^*i.v. with concurrently with i.n. LPS (A). Alveolar permeability assessed by BAL albumin or total protein (B, C), neutrophil recruitment (D), and BAL IL-6 (E) shown 72 hours after i.n. LPS. BAL protein measurements expressed relative to the mean of unoperated control *Ccr2^wt^* transfer for each cohort. n = 3-6 per group, 3 cohorts. Mean ± SEM, Welch’s t-test p-value shown. * p < 0.05, ** p < 0.01. CLP, Cecal Ligation and Puncture. LPS, Lipopolysaccharide. BAL, bronchoalveolar lavage. PMN, polymorphonuclear.

### Post-CLP Ly6C^hi^ monocytes promote neutrophil activation and degranulation in response to LPS

Lung permeability in LPS-induced lung injury is neutrophil-dependent^40–44^. However, despite alterations in lung permeability, we observed no differences in neutrophil recruitment after monocyte depletion and adoptive transfer (Figures 1E, 2D), suggesting that the mechanism by which post-CLP monocytes enhance lung injury is not via augmented neutrophil recruitment. Therefore, we hypothesized that post-CLP monocytes may enhance neutrophil activation. We previously observed elevated levels of S100A8/A9, a damage associated-molecular pattern protein released upon neutrophil activation^45–48^, in the BAL of post-CLP mice following i.n. LPS^12^. Here, we found that adoptive transfer of post-CLP monocytes increased (Figure 3A), whereas monocyte depletion reduced, BAL S100A8/A9 in post-CLP mice (Figure 3B), suggesting that post-CLP monocytes activate neutrophils.

**Figure 3.**
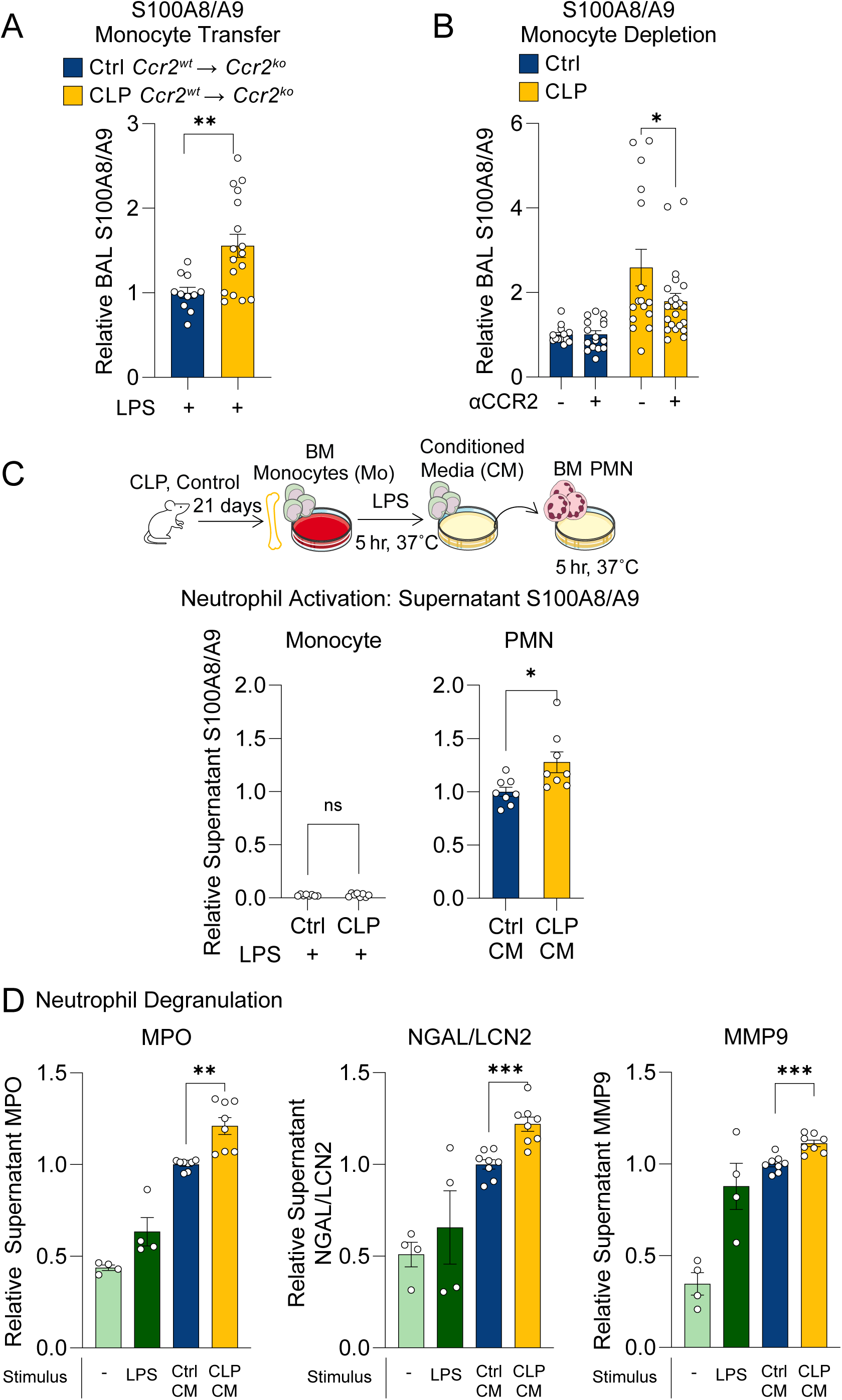
Post-CLP monocytes promote activation and degranulation of neutrophils. BAL S100A8/A9 following adoptive transfer (A) and antibody-mediated depletion of bone marrow Ly6C^hi^ monocytes (B). Monocyte and neutrophil S100A8/A9 production in supernatants following stimulation with LPS or monocyte conditioned media (CM), respectively (C). Neutrophil granule proteins were measured to assess degranulation induced by monocyte CM (D). n = 4 mice per group, 2 cohorts. Supernatant protein expressed relative to the mean of unoperated control monocyte CM. Mean ± SEM, Welch’s t-test p-value shown. * p < 0.05, ** p < 0.01, *** p <0.01. CLP, Cecal Ligation and Puncture. LPS, Lipopolysaccharide. BAL, bronchoalveolar lavage. MPO, Myeloperoxidase. NGAL/LCN2, neutrophil gelatinase-associated lipocalin. MMP9, matrix metallopeptidase-9.

A key facet of neutrophil activation is degranulation, the release of granule contents — such as proteolytic enzymes, antimicrobial peptides, and other substances that contribute to lung injury ^49–52^ — into the extracellular space. To investigate whether post-CLP monocytes directly induce neutrophil activation and degranulation, we isolated BM Ly6C^hi^ monocytes from post-CLP mice, stimulated with LPS, and transferred the monocyte conditioned media (CM) to naïve BM neutrophils (Figure 3C). The CM from post-CLP monocytes induced greater release of S100A8/A9 (Figure 3D) and neutrophil granule proteins (myeloperoxidase (MPO), neutrophil gelatinase-associated lipocalin (NGAL), matrix metallopeptidase 9 (MMP9)) than CM from unoperated control monocytes (Figure 3D). These data suggest that post-CLP monocytes enhance neutrophil activation and degranulation, thereby contributing to enhanced lung injury.

### Enhanced lung injury in post-CLP mice persists for three months

Prior infection or inflammation can have durable effects on subsequent inflammatory responses^22–26^. Therefore, we hypothesized that the enhanced lung injury response observed in sepsis survivor mice may be durable far beyond 3-wk post-CLP. We therefore evaluated the lung injury response at 3 months (3-mo) post-CLP (Figure 4A). Remarkably, post-CLP mice exhibited significantly increased lung permeability and epithelial injury compared to control mice (Figure 4B, 4C). This was accompanied by increased airspace neutrophils and IL-6 levels (Figure 4D, 4E). Adoptive transfer of monocytes from 3-mo post-CLP mice resulted in increased lung permeability relative to transfer of unoperated control monocytes (Figure 4F). These findings confirm that enhanced lung injury following recovery from sepsis is a durable phenotype that is mediated by monocytes. This durable pro-inflammatory phenotype is consistent with reprogramming of monocytes, wherein monocytes are primed for enhanced responses to a secondary stimulus.

**Figure 4.**
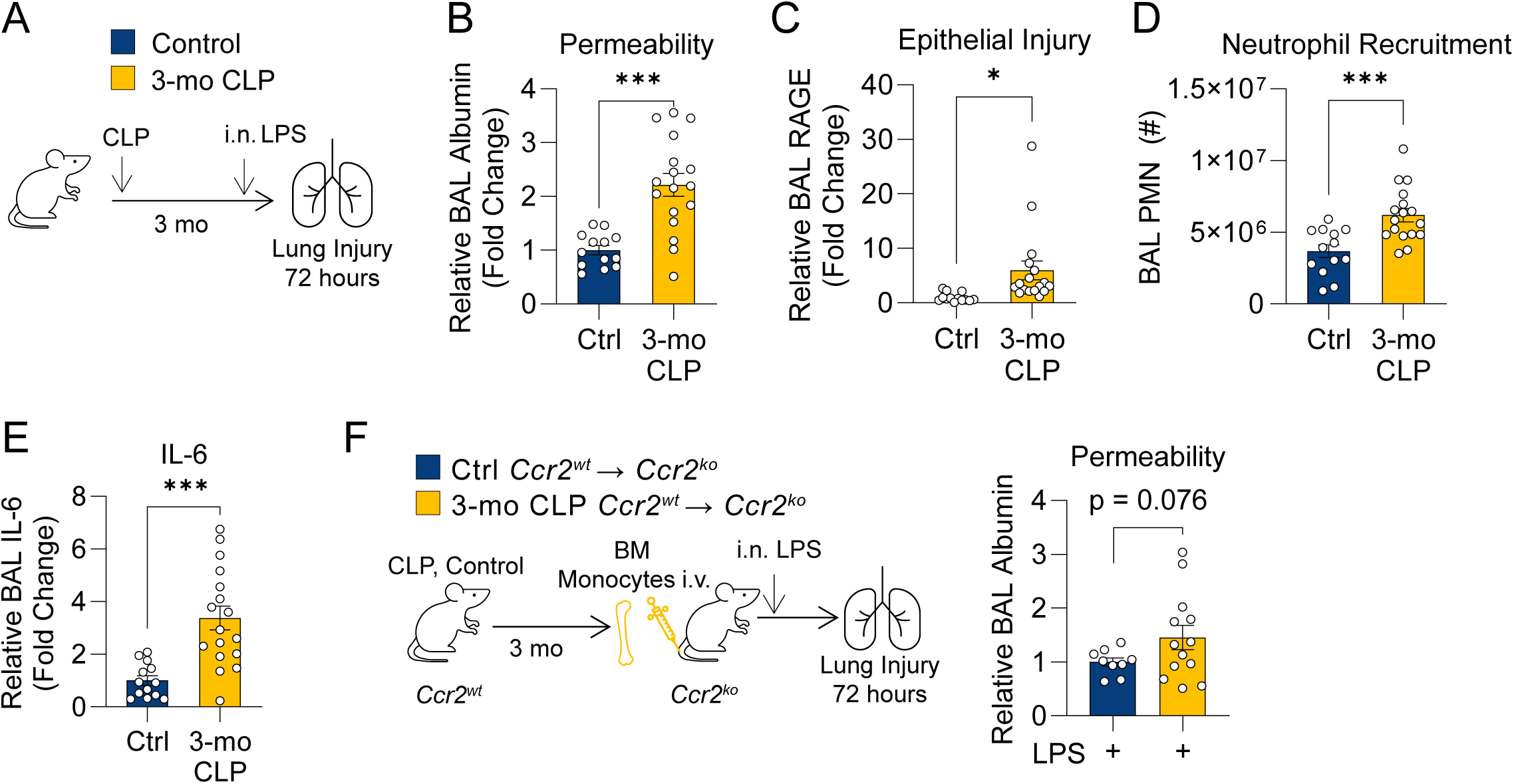
Monocyte mediated enhanced lung injury in post-CLP mice is durable. 3-mo post-CLP and age-matched control mice were administered i.n. LPS (A). Alveolar permeability assessed by BAL albumin (B), epithelial injury assessed by BAL RAGE (C), neutrophil recruitment (D), and BAL IL-6 (E) measured 72-hr after i.n. LPS. Bone marrow Ly6C^hi^ monocytes were isolated from *Ccr2^wt^* age-matched unoperated control, and *Ccr2^wt^*3-mo post-CLP mice and administered into *Ccr2^-/-^* mice i.v. concurrently with i.n. LPS, alveolar permeability assessed by BAL albumin (F). n = 5-10 per group, 2 cohorts (A-E), n = 2-6 per group, 3 cohorts (F). Mean ± SEM, Welch’s t-test p-value shown. * p < 0.05, ** p < 0.01, *** p < 0.001. CLP, Cecal Ligation and Puncture. LPS, Lipopolysaccharide. BAL, bronchoalveolar lavage. RAGE, receptor for advanced glycation end-products. PMN, polymorphonuclear. BM, bone marrow.

### CLP elicits pro-inflammatory transcriptional and epigenetic reprogramming of Ly6C^hi^ monocytes

The enhanced lung injury and neutrophil activation mediated by monocytes after sepsis, particularly in contrast to the protective role of monocytes in lung injury without preceding sepsis, imply that sepsis reprograms Ly6C^hi^ monocytes toward a primed state that promotes a more robust pro-inflammatory response to secondary stimuli. Such immune reprogramming following an initial stimulus is typically attributable to transcriptional and epigenetic alterations^23^. We hypothesized that CLP induces persistent alterations in the transcriptome and/or epigenome of monocytes. To investigate these alterations, we performed bulk RNA- and ATAC-sequencing on mature BM Ly6C^hi^ monocytes isolated at 3-wk post-CLP (Figure S5). Pathway analysis of gene expression data revealed that post-CLP monocytes were enriched for pathways related to HIF-1α, cell cycle, glucose/lipid metabolism, and inflammatory signaling relative to control (Figure 5A). Specifically, post-CLP monocytes upregulated TLR (*Tlr4*, *Lbp, Myd88*, *Mapk13*) and JAK-STAT pathway genes while downregulating other anti-inflammatory genes (*Nfkbia, Nfkbie*, *Igf1*) (Figure 5B). ATAC-seq identified enrichment of uniquely accessible promoter and enhancer regions, indicating active or poised transcription, in post-CLP monocytes. Accessible promoters were linked to genes in glycolytic and inflammatory pathways (Figure 5C). The JAK-STAT signaling pathway was enriched in both the promoter (Figure 5C) and enhancer (Figure 5D) regions in post-CLP monocytes. In contrast, promoter regions in monocyte from unoperated control mice were enriched for homeostatic pathways, including vesicle transport, fatty acid biosynthesis/signaling, acetyl-CoA handling, and nucleotide metabolism (Figure S6). To identify the transcription factors (TF) that potentially binding these open genomic regions, we conducted motif enrichment analysis^53^. This identified Fos:Jun motifs for the dimeric Activator Protein-1 (AP-1) TF in enhancer regions of post-CLP monocytes (Figure 5E), including a nearly 3-fold increase in the number of genes with at least one enhancer peak containing the AP-1 motif (Figure 4F). In other inflammatory contexts, AP-1 binding to enhancer regions of DNA is essential for maintaining chromatin accessibility and the enhanced inflammatory mediator production in response to secondary challenge that characterizes a durable form of reprogramming known as immune memory^54–56^. Together, these data demonstrate transcriptional and epigenetic reprogramming of BM Ly6C^hi^ monocytes towards a persistently primed state.

**Figure 5.**
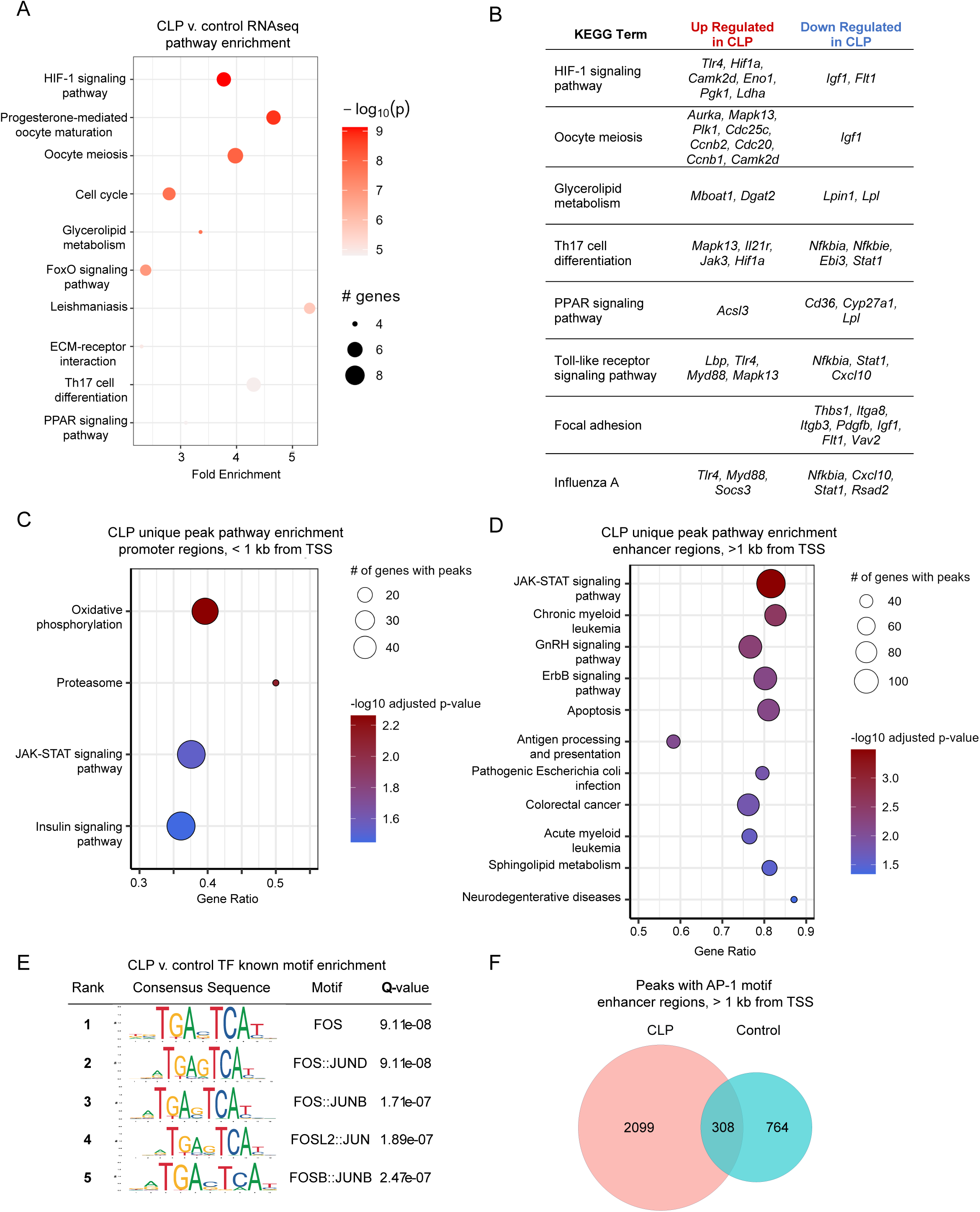
Transcriptional and epigenetic pathways contributing to post-CLP monocyte priming. Ly6C^hi^ monocytes were isolated from the bone marrow for transcriptomic and epigenomic profiling. Top 10 transcriptomic pathways enriched in 3-wk post-CLP monocytes relative to control (A). Selected transcriptomic pathways enriched in post-CLP monocytes with upregulated and downregulated genes per pathway (B). In a separate experiment, ATAC-seq was performed, unique peaks were identified, and pathway enrichment performed on promoter (C) and enhancer regions (D). Known motif enrichment performed relative to control monocytes genomic background (E). Quantification of peaks containing the AP-1 binding motif in post-CLP and control monocytes (F). n = 4-5 mice per group. CLP, Cecal Ligation and Puncture. TSS, transcription start site.

### Sepsis induced by CLP reprograms monocytes toward the neutrophil-like lineage

Recent scRNAseq studies have begun to uncover subsets of classical monocytes (Figure 6A), which are derived from different BM progenitors and express typical genes for neutrophils and DC, neutrophil-like and DC-like monocytes, respectively ^33–36,57,58^. Deeper interrogation of the multiomics data revealed that post-CLP monocytes upregulated neutrophil-specific genes (e.g., neutrophil granule protein, *Ngp*, proteinase-3 *Prtn3*, lipocalin-2 *Lcn2*), while downregulating dendritic cell (DC)-specific genes (e.g., *Cd209a*) (Figure 6B). Although the transcriptomes of these subsets have been characterized by scRNAseq, to our knowledge, aside from isolated studies ^38,59^, the specific functions of these novel monocyte subsets remain unexplored.

**Figure 6.**
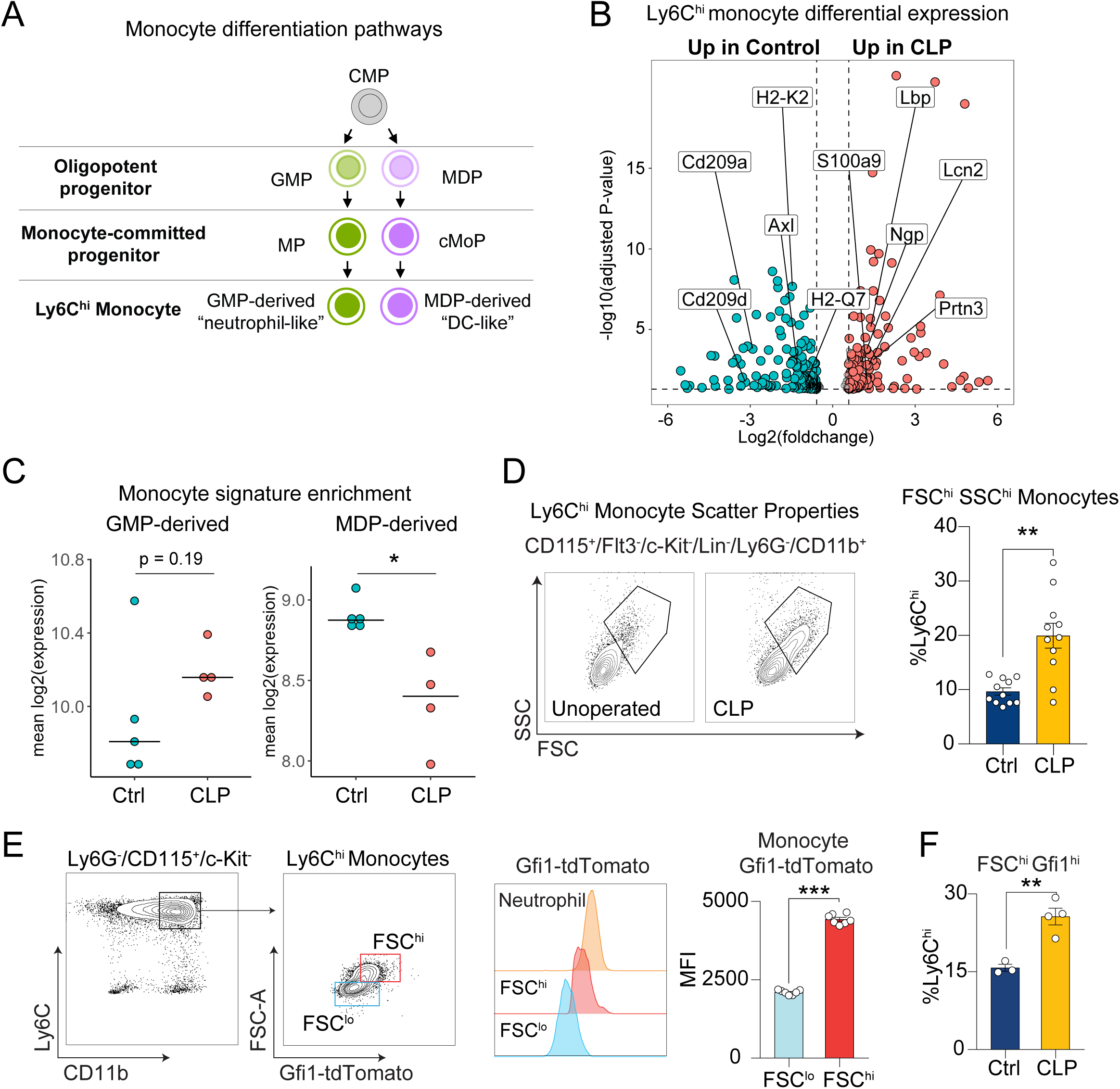
Neutrophil-like monocytes are expanded in post-CLP mice. Monocyte differentiation pathways^33^ (A). Differential expression in Ly6C^hi^ monocytes highlighting neutrophil- and DC-specific genes (B). Enrichment of GMP-derived and MDP-derived monocyte signatures^33^ in Ly6C^hi^ monocytes (C). Scatter properties of Ly6C^hi^ monocytes in post-CLP and unoperated control (D). Gfi1-tdTomato fluorescent intensity in Neutrophils and FSC^hi^ or FSC^lo^ monocytes (E). Proportions of FSC^hi^Gfi1-tdTomato Ly6C^hi^ monocytes in post-CLP and unoperated control mice (F). n= 4-5 per group, 2 cohorts (D), n=3-5 mice per group, 1 cohort (B, C, E, F). Mean, Mean ± SEM and Welch’s t-test (D-F) are shown. * p < 0.05, ** p < 0.01.. MDP, monocyte-dendritic cell progenitor. cMoP, common monocyte progenitor. GMP, granulocyte-monocyte progenitor. MP, monocyte progenitor. CLP, cecal ligation and puncture. FSC, forward scatter. SSC, side scatter.

Moreover, to our knowledge, these monocyte subsets have not been characterized in the context of prior sepsis and lung injury. Based on the increased expression of neutrophil-like monocyte genes in post-CLP monocytes (Figure 6B), we hypothesized that CLP reprograms monocytes towards a neutrophil-like state. To determine whether CLP reprograms monocytes toward a neutrophil-like state, we conducted gene signature enrichment analysis using gene modules specific to neutrophil-like and DC-like monocytes^33^. Monocytes from post-CLP mice were enriched for the neutrophil-like monocyte signature, while control monocytes were enriched for the DC-like monocyte signature, suggesting that CLP induces expansion of the neutrophil-like monocyte population (Figure 6C).

To confirm expansion of the neutrophil-like monocyte population following CLP, we analyzed forward-scatter (FSC) and side-scatter (SSC) flow cytometric properties of Ly6C^hi^ monocytes, as larger size and increased granularity have been shown to identify neutrophil-like monocytes^33^. We identified two populations of monocytes within the Ly6C^hi^ fraction (Figure S5): FSC^lo^SSC^lo^ and FSC^hi^SSC^hi^ (Figure 6D). Post-CLP mice exhibited a 2-3 fold increase in FSC^hi^SSC^hi^ Ly6C^hi^ cells, consistent with neutrophil-like monocyte expansion. To further validate expansion of neutrophil-like monocytes after CLP, we used *Gfi1*-tdTomato mice, which identify this lineage due to their higher levels of *Gfi1*^33^. We observed higher *Gfi1* expression in the Ly6C^hi^ monocytes (Figure 6E). In fact, tomato fluorescence in the FSC^hi^ population was comparable to neutrophils and significantly greater than FSC^lo^ monocytes, strongly suggesting that the Gfi1^hi^ cells represent neutrophil-like monocytes. To confirm that sepsis induces expansion of neutrophil-like monocytes, we performed CLP on *Gfi1*-tdTomato mice and found that Gfi1^hi^FSC^hi^ monocytes were significantly increased in post-CLP mice (Figure 6F). These data confirm that CLP reprograms monocytes toward the neutrophil-like lineage.

### CLP reprograms bone marrow monocyte progenitors toward the neutrophil-like lineage

Given that sepsis leads to a durable expansion of the neutrophil-like monocyte population, but the half-life of monocytes is only hours to days^60,61^, we hypothesized that sepsis survival shifts hematopoiesis and monocyte fate toward the neutrophil-like lineage at the level of monocyte progenitors. Previous studies using ex vivo culture of monocyte progenitors revealed that neutrophil-like monocytes arise from the granulocyte-monocyte progenitor (GMP) via their own monocyte-committed progenitor (MP), while dendritic cell (DC)-like monocytes arise from the monocyte-DC progenitor (MDP) via the common monocyte progenitor (cMoP) (Figure 6A). To specifically assess expansion of the GMP-derived lineage, we flow sorted the MP+cMoP fraction (Figure S7), performed bulk RNA-sequencing, and assessed for enrichment of MP and cMoP progenitor gene signatures. Consistent with our hypothesis, post-CLP BM progenitors were enriched for the MP signature (Figure 7A). We then directly assessed for GMP expansion in the BM of post-CLP mice, using established surface markers of GMPs (Figure S7)^62^. We demonstrated an increase in GMP progenitors in post-CLP mice compared to controls, without changes in MDPs (Figure 7B). Consistent with expanded GMP, which produce neutrophils as well as monocytes, we found a concomitant increase in white blood cells characterized by increased circulating neutrophils and monocytes in post-CLP mice (Figure 7C).

**Figure 7.**
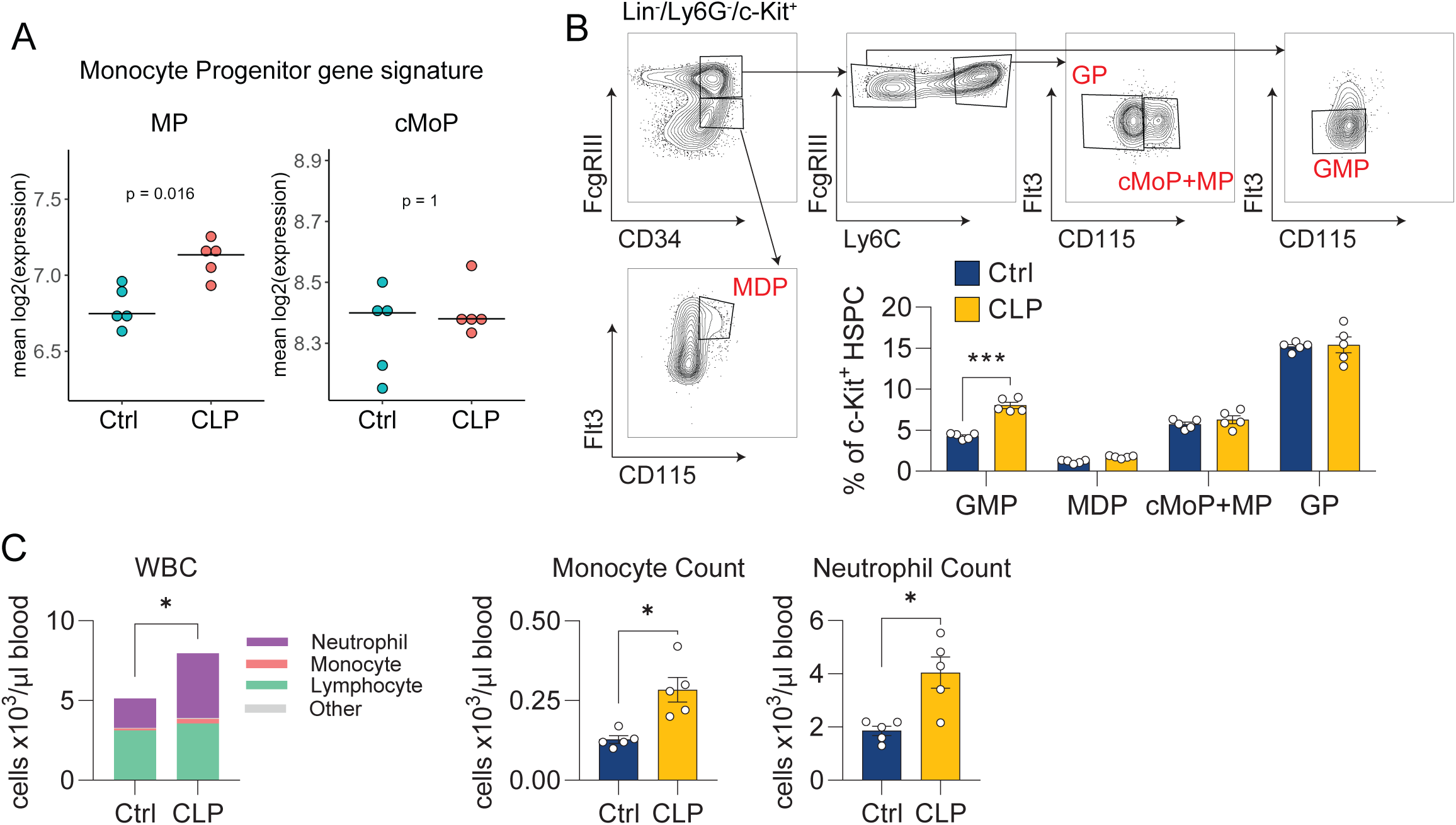
CLP reprograms progenitors toward GMP-lineage. Monocyte committed BM progenitors (MP+cMoP enriched) were isolated from BM and tested for enrichment of MP (GMP-derived) and cMoP (MDP-derived) signatures (A)^33^. Quantification of myeloid progenitors by flow cytometry (B). Peripheral blood counts (C). n = 4-5 per group, 1 cohort (A-C). Mean, Wilcoxon rank-sum (A), Mean ± SEM and Welch’s t-test (B, C) are shown. * p < 0.05, ** p < 0.01, *** p <0.01. MDP, monocyte-dendritic cell progenitor. cMoP, common monocyte progenitor. GMP, granulocyte-monocyte progenitor. MP, monocyte progenitor. CLP, cecal ligation and puncture. WBC, white blood cell.

In summary, these data suggest that sepsis induced by CLP reprograms hematopoietic progenitors resulting in a shift in monocyte ontogeny toward the GMP lineage, thereby expanding neutrophil-like monocytes and their progenitors, MPs and GMPs. Coupled with the demonstrated functional role of post-CLP monocytes in exacerbating LPS-induced lung injury (Figure 1,2), these data suggest that the a shift in BM ontogeny towards the neutrophil-like monocyte subset promotes enhanced lung injury in post-CLP mice.

### Neutrophil-like monocytes are expanded by acute human sepsis and monocyte counts predict long-term mortality in human sepsis survivors

Patients who survive sepsis are at increased risk of long-term rehospitalization and death^2,3^, yet the underlying mechanisms remain unclear. Given the role of monocytes in mediating enhanced acute lung injury in sepsis survivor mice (Figure 1, 2), we hypothesized that mobilization of monocytes may similarly predict poor long-term outcomes after sepsis in humans. We previously demonstrated that abnormal leukocyte counts at the time of hospital discharge predict 90-day rehospitalization and/or death^9^ in patients surviving sepsis; here, we examined whether absolute monocyte count (AMC) at hospital discharge would also predict 90-day mortality. Using a United States Veteran’s Affairs (VA) health system dataset^63^, we identified 92,165 patients that met inclusion criteria for sepsis (Figure S8), 13.6% of whom died within 90-days (Supplemental Table 1), consistent with our prior findings^9^. We found that absolute monocyte count (AMC) was associated with an increased hazard ratio for 90-day mortality (Figure 8A).

**Figure 8.**
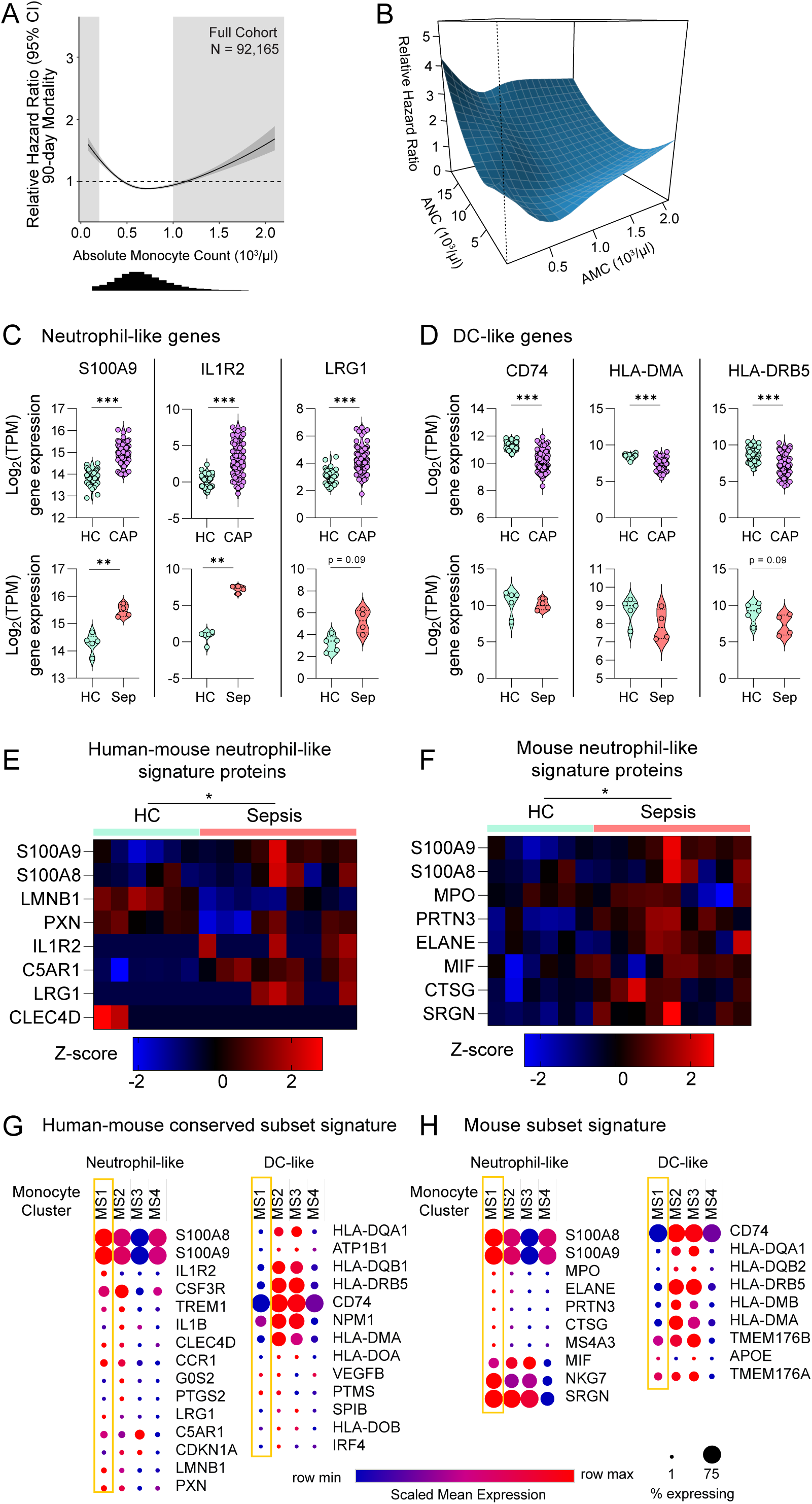
Neutrophil-like monocyte lineage in patients with sepsis. Unadjusted Cox proportional hazard regression (95% CI, dark gray) with relative hazard ratio for 90-day mortality for absolute monocyte count (AMC), histogram of AMC distribution shown below (A). Light gray boxes indicate values outside of the normal clinical range. Three-dimensional interaction surface showing the relationship of AMC and absolute neutrophil count (ANC) with 90-day mortality (B). Selected neutrophil-like and DC-like gene expression in CD14+ monocytes in patients with acute sepsis (n = 4 sepsis, 5 healthy)^65^ and community acquired pneumonia (n = 69 CAP, 41 healthy) (C, D)^66^. Conserved human-mouse^68^ (E) and mouse-specific^36^ (F) neutrophil-like signature proteins in classical monocytes from patients with sepsis^67^. Proteins detected in more than 1 sample across conditions were included in the heatmap, missing values were imputed as described in the methods. Conserved human-mouse (G) or mouse-specific signatures (H) across monocyte substates (MS1-4) in scRNAseq performed in patients with acute bacterial infection^64^. Mean, 25^th^ and 75^th^ percentile shown (C, D). Multivariate-ANOVA p-value shown (E, F). * p < 0.05, ** p < 0.01, *** p < 0.001. HC, healthy control. Sep, sepsis. CAP, community acquired pneumonia.

We next investigated whether the monocytes associated with mortality in sepsis patients were enriched for neutrophil-like monocytes. In our murine sepsis model, concurrent expansion of neutrophils and monocytes (Figure 7C) reflected mobilization of the GMP-lineage. We have shown that AMC (Figure 8A) and absolute neutrophil counts (ANC) were associated with 90-day mortality^9^. To gain insight into whether the associations between increased AMC and mortality and increased ANC and mortality may indicate that mobilization of a common progenitor is linked to poor outcomes, we first analyzed whether the predictive value of monocyte count was dependent on neutrophil count by plotting the hazard ratio surface for the interaction of AMC and ANC for 90-day mortality (Figure 8B). The hazard ratio for AMC increased most markedly as ANC increased. This indicates that the prognostic value of AMC is highly dependent on the concomitant ANC, suggesting that AMCs are most predictive of poor outcome when the common progenitor is mobilized. Since monocytes and neutrophils both derive from GMPs, our findings suggest persistent mobilization of the GMP-lineage is highly associated with poor outcome in human sepsis patients.

To acquire further evidence to establish whether neutrophil-like monocytes were expanded in acute human sepsis, we interrogated publicly available RNAseq datasets of bulk human CD14+ monocytes from patients with sepsis for evidence of a neutrophil-like gene signature^64–67^. To assess enrichment of monocyte subsets, we utilized neutrophil-like and DC-like monocyte gene signatures conserved between mice and humans^68^. As expected, these signatures detected the enrichment of GMP-derived neutrophil-like monocytes in our CLP mouse model of sepsis (Figure S9). We then assessed the expression of individual neutrophil-like and DC-like genes by CD14+ monocytes using bulk RNAseq data from multiple distinct cohorts of patients with sepsis or pneumonia ^65,66^. We observed upregulation of conserved neutrophil-like genes (*S100A9*, *IL1R2*, *LRG1*) and downregulation of genes associated with DCs (*CD74*, *HLA-DMA*, *HLA-DRB5*) (Figure 8C, 8D). To validate the expansion of neutrophil-like monocytes in human patients, we next examined the proteome of CD14^+^ monocytes in patients with bacterial sepsis^67^ (Figure 8E, 8F). We again found enrichment of neutrophil-like proteins, in the monocytes of patients with sepsis. This consistent observation of increased levels of neutrophil genes and proteins across multiple independent cohorts of acute pneumonia and sepsis suggests a shift toward a neutrophil-like state in acute infectious/inflammatory human disease of diverse etiologies.

Finally, to confirm that enrichment of the neutrophil-like signature in bulk CD14^+^ monocytes reflects expansion of a neutrophil-like population in acute sepsis, we explored a single cell RNA-sequencing (scRNA-seq) dataset of CD14+ monocytes from patients with acute bacterial infection and sepsis^64^. Unbiased clustering revealed four monocyte states (MS1-MS4)^64^; the MS1 cluster was previously found to be expanded in patients with bacterial infection and sepsis. To determine whether MS1 represents a neutrophil-like monocyte, we examined the conserved monocyte subset gene signatures across the four monocyte states. Consistent with our hypothesis, the MS1 cluster consistently upregulated neutrophil-like and downregulated DC-like genes, suggesting that MS1 is indeed a neutrophil-like state (Figure 8B). Thus, the alterations observed in bulk monocytes from septic patients (Figure 8C-F) likely reflect expansion of neutrophil-like monocytes. Taken together, these data strongly suggest that neutrophil-like monocytes are mobilized in both mice and humans with sepsis and promote poor outcomes.

## Discussion

As survival rates for sepsis improve, more patients are at risk of post-sepsis complications, including rehospitalization with new organ injury. Herein, we demonstrated that sepsis induced by CLP in mice durably reprograms Ly6C^hi^ monocytes, which predisposes to enhanced lung injury upon LPS challenge. We identified key transcriptomic and epigenomic signatures persistently activated after CLP, including JAK-STAT and AP-1. Moreover, CLP induces Ly6C^hi^ monocytes and their progenitors to shift toward the GMP-derived neutrophil-like lineage. Importantly, we discovered that monocyte counts predict long-term outcomes in human sepsis survivors and these monocytes are similarly enriched for a neutrophil-like signature. This suggests that specific monocyte subsets with distinct BM origins influence the trajectory of lung injury. Monocytes reprogrammed by sepsis also promote neutrophil activation. These data support a conceptual model whereby sepsis reprograms the immune activation state and ontogeny of monocytes and their progenitors leading to a sustained risk for new organ injury.

These findings significantly advance our understanding of functional monocyte heterogeneity. Although an emerging literature has revealed that Ly6C^hi^ monocytes are heterogenous^33–36^, the specific contexts in which specific monocyte subsets arise and their functional roles in disease pathogenesis remain significant and clinically relevant gaps in our knowledge^58,69^. Neutrophil-like and DC-like subsets of Ly6C^hi^ monocytes have recently been identified by scRNAseq in both homeostasis and sterile inflammation in multiple species^33–36^. Here, we demonstrate that neutrophil-like monocytes expand in murine and human sepsis. Using the CLP mouse model, we demonstrate expansion of neutrophil-like monocytes via multiple methods: gene expression analysis, flow cytometric (FSC/SSC) characteristics, and the use of a Gfi1-reporter mouse (Figure 6, 7). Supporting our findings, monocytes that appear transcriptionally similar were recently reported to emerge in the spleen after CLP^70^, although they were not specifically characterized as GMP-lineage neutrophil-like monocytes. Importantly, we also confirm the expansion of neutrophil-like monocytes in humans with sepsis as well as pneumonia (Figure 8), demonstrating that this phenomenon is conserved across species and across multiple acute infectious/inflammatory human diseases.

However, the greatest novelty and impact of this work lie in the functional studies. Our data strongly suggest that the durable expansion of the neutrophil-like monocytes induced by sepsis (Figure 6) exacerbates subsequent lung injury in mice (Figures 1, 2, 4). Following isolated reports^38,59^, this represents one of the first demonstrations of a specific functional role of a monocyte subset. Moreover, for the first time to our knowledge, our data implicate neutrophil-like monocytes as functionally involved in human disease pathogenesis. Although the MS1 gene signature was shown to be associated with bacterial infection^64^ and risk of ARDS in severe sepsis^71^, causality was not established, nor were MS1 monocytes previously characterized as neutrophil-like. Here, we demonstrate that the human MS1 monocyte population is analogous to the GMP-derived neutrophil-like monocytes in mice (Figure 8). By demonstrating that murine neutrophil-like monocytes recapitulate the human MS1 monocytes and that murine neutrophil-like monocytes are pathogenic, our data strongly suggest that clinical association of the MS1 signature with poor outcomes reflects an underlying pathogenic role of the MS1 monocytes in human disease.

An additional novel key finding is that neutrophil-like monocytes can be pathogenic, not just protective. Previous functional studies, though sparse, have demonstrated a protective role for neutrophil-like monocytes ^34,38,59^. Ly6C^hi^ monocytes expressing high levels of neutrophil granule genes and *Ym1* (chitinase-like 3, *Chil3*) are protective against experimental colitis^59^ and brain injury^38^. In contrast, we show that post-sepsis murine neutrophil-like monocytes exacerbate LPS-induced acute lung injury, and a similar neutrophil-like transcriptional signature is associated with poor outcomes in human sepsis^71^. Interestingly, this contrasts with monocytes in acute lung injury without preceding sepsis, which are protective (Figure 1, 2, 4)^14^. Our results suggest that post-sepsis neutrophil-like monocytes may adopt an alternate inflammatory program without the protective features seen in colitis, brain injury, and lung injury without prior sepsis. This implies that the functions of neutrophil-like monocytes in inflammatory organ injury may depend significantly on disease or tissue context. Additionally, it raises the possibility of further heterogeneity among neutrophil-like monocytes, with those identified in earlier studies differing from the subsets we have observed. By uncovering previously unappreciated complexity in monocyte heterogeneity, our study underscores the nascent state of this field and the need for new avenues of investigation.

These findings fundamentally reshape our understanding of immune memory, demonstrating that it may occur both through epigenetic priming of proinflammatory genes and through the reprogramming of hematopoietic progenitors toward alternate monocyte lineages. Innate immune memory, the durable reprogramming of immune progenitors for enhanced responses to secondary stimuli^55,72,73^, may protect the host from secondary bacterial infections^72–74^ but can be maladaptive in other inflammatory diseases^72,75–77^. Immune memory is established through alterations in chromatin accessibility at proinflammatory genes, induced by a primary stimulus, thereby facilitating transcription factor binding upon a secondary insult^25^. Accordingly, we found that sepsis alters chromatin accessibility with enrichment of AP-1 motifs and enhanced JAK-STAT, HIF-1α and TLR signaling - which are established mechanisms of immune memory^10,54,55,73,78,79^ - in association with an enhanced lung injury phenotype that persists for at least 3 months. Thus, our findings suggest that innate immune memory, via classic mechanisms, leads to monocyte pathogenicity in lung injury after sepsis. We demonstrate that sepsis alters lineage specification leading to expansion of GMPs and neutrophil-like monocytes. While prior literature hints that neutrophil-like monocyte expansion might be an aspect of innate immune memory^57,79^, our study provides substantial evidence for this phenomenon. Previous reports of immune memory have noted a myeloid differentiation bias in hematopoietic progenitors^55,72,73^: our findings suggest this may reflect mobilization of GMP lineage monocytes. Trained immunity induced by BCG vaccination in humans is associated with expansion of a neutrophil-like monocyte subset^79^. Similarly, in a primate model of chronic alcohol exposure after weeks-months of abstinence, monocytes exhibit an enhanced response to secondary stimuli and hematopoietic progenitors are biased toward production of neutrophil-like monocytes^57^. Our paper further advances the field by demonstrating these shifts in ontogeny in humans survivors of sepsis, as evidenced by a neutrophil-like gene signature during acute sepsis and correlations of elevated monocyte and neutrophil counts with mortality (Figures 6 and 8). This seminal discovery that immune memory may involve both altered inflammatory programming and shifts in ontogeny in hematopoietic progenitors establishes a foundation for future studies to assess the role of altered ontogeny in immune memory in other contexts, the underlying mechanisms, and how shifts in ontogeny and altered inflammatory activation status may interact to regulate monocyte function.

Finally, this study offers significant insights into the pathogenesis of acute lung injury. First, while inflammatory reprogramming during acute sepsis has been linked to adverse short-term clinical outcomes^26^, our research elucidates the effects of durable reprogramming after sepsis and its contribution to poor long-term outcomes, including new organ injury. In mice, sepsis durably reprogrammed Ly6C^hi^ monocytes, resulting in increased lung injury in response to subsequent LPS exposure for at least 3 months. Second, these findings transform our understanding of monocyte functional heterogeneity in acute lung injury. While diverse functions of monocytes in different etiologies of ALI^14–17,19–21,80–82^ have supported the prevailing paradigm that their activation status and function are dictated by the local lung milieu^13^, we demonstrate here that functional monocyte heterogeneity may also be shaped by the imprinting of monocyte progenitors in the BM leading to the expansion of specific monocyte subsets. The interplay between ontogeny and the local lung environment in shaping monocyte function across diverse etiologies of ALI is an important topic for future research. Third, the mechanisms through which immune memory contributes to organ injury are poorly understood. We show a mechanism through which monocyte reprogramming exacerbates lung injury. Neutrophils are pathogenic in lung injury and the acute respiratory distress syndrome (ARDS)^40–44^. While neutrophil activation may be adaptive to calibrate host defense against future bacterial infections, proteases and myeloperoxidase released upon degranulation have detrimental effects on the alveolar capillary barrier^45–48^. Here we show that post-CLP monocytes enhance neutrophil activation, including degranulation, in the absence of effects on recruitment, suggesting that maladaptive monocyte programming promotes lung injury via neutrophil activation. Future investigation into the role of monocyte-neutrophil crosstalk in lung injury is warranted. Finally, these mechanisms of post-sepsis lung injury hold significant therapeutic implications. Additional studies on monocyte subset variation in the BM and their functions in health and disease may enable us to predict and/or prevent the development of organ injury. Targeting both immune cell activation (e.g, AP-1 or JAK-STAT signaling) and ontogeny, as well as monocyte-neutrophil crosstalk, may have therapeutic utility in preventing pulmonary complications in sepsis survivors.

This study has some limitations. While we utilized LPS to induce lung injury, future research on how post-sepsis monocyte reprogramming affects lung injury and host defense in the setting of live bacterial or viral infection is warranted. Although we demonstrate that elevated monocyte counts, enriched for neutrophil-like monocytes, predict poor clinical outcomes in patients with acute infection, further studies need to evaluate specific monocyte subsets in the circulation and progenitor populations in the BM of patients who have survived sepsis. This will be crucial to completely understand durable immune reprogramming, shifts in ontogeny, and their long-term clinical consequences in human sepsis. Finally, we must develop methods (e.g., Cre drivers) to specifically target neutrophil-like monocytes to definitively establish their pathogenicity.

In summary, our findings provide insight into fundamental questions in the emerging field of monocyte heterogeneity^58^: the extent to which Ly6C^hi^ monocyte subsets and inflammatory context regulate functional heterogeneity in health and disease. Our data suggest that Ly6C^hi^ monocytes enhance the lung injury response to LPS by promoting neutrophil degranulation. Moreover, we observe for the first time a functional role for neutrophil-like monocytes in the pathogenesis of acute lung injury. With the discovery of a potentially subset-specific phenotype, these observations establish a foundation for future investigations to address key gaps in our understanding of how monocyte subset and immune memory might collaborate to enhance tissue injury responses. The functional role of specific monocyte subsets in the BM in lung injury, their selection by the lung inflammatory milieu, and their effect on the phenotype of recruited macrophages will help refine our prior understanding of monocyte/macrophage mediated inflammation in the lungs. Finally, the persistent nature of hematopoietic reprogramming after sepsis in mice identifies a potential therapeutic target for the prevention of post-sepsis organ injury leading to rehospitalization in humans.

## Materials and Methods

### Mouse studies

Male 8–12 weeks old C57BL/6J (strain 000664) and *Ccr2^-/-^*(strain 004999) mice were purchased from Jackson Laboratories. All procedures involving animals were undertaken in strict accordance with the recommendations of the Guide for the Care and Use of Laboratory Animals by the National Institutes of Health. The University of Michigan facility maintains a Specific Pathogen Free barrier environment.

### Cecal ligation and puncture

CLP was performed as previously described^12,83^. Briefly, animal suffering and distress were minimized using analgesia with local lidocaine, as well as anesthesia with ketamine and xylazine. Under aseptic conditions, a 1–2 cm laparotomy was performed. The cecum was ligated with a silk suture and punctured once through-and-through with a 19-gauge needle. The incision was closed with surgical clips. Imipenem/cilastatin (Merck, 0.5 mg/mouse in 200 µl of normal saline) and normal saline (0.5 mL) were administered subcutaneously to all CLP mice immediately following surgery. This method induces polymicrobial bacterial peritonitis with disseminated infection, an average mortality of 10%, and clearance of bacterial cultures within 5-7 days^12,83,84^. By three weeks, there is no difference in weight between groups ^12^. We also did not observe any additional mortality during the three week to three month recovery period, and aerobic peritoneal cultures in 3-mo post-CLP mice remained negative (data not shown).

### Lipopolysaccharide-induced lung injury

*Escherichia coli* LPS (O111:B4, Sigma) was administered i.n. at predetermined time points (18 to 84 days) after CLP or in age-matched unoperated control mice. We utilized unoperated control mice because we previously established that sham surgery and unoperated control mice have similar lung injury responses to LPS at 3 weeks post-surgery^12^. Briefly, mice were anesthetized using ketamine and xylazine. LPS dissolved in sterile normal saline (50 µg, 1 µg/µl) was administered 25 µl per nostril. Mice were euthanized 24 or 72 hours after LPS administration. The University of Michigan policy for humane endpoints was followed.

### Monocyte depletion

Monocyte depletion was performed using an anti-CCR2 antibody (clone MC-21^85^) or rat IgG2b isotype (clone LTF-2, BioXCel or clone MC-67) administered intraperitoneally (i.p., 40 μg) four days prior (day -4), two days prior (day -2), and at the time of LPS administration (day 0). Treatments were randomized within cage. At 72 hours after LPS, post-CLP mice treated with anti-CCR2 antibody had less mononuclear cells (DiffQuik) in the BAL than isotype treated mice, indicating monocyte depletion. One cohort (out of 4 total depletions) showed instead elevated mononuclear cell counts in the BAL relative to isotype treated control for all mice, this was inconsistent with monocyte depletion and was excluded from analysis.

### BM isolation and immunomagnetic monocyte isolation

BM was isolated from femurs and tibia of post-CLP mice (days 21 and 84) and age-matched control mice. Briefly, mice were euthanized with ketamine and xylazine. In a biological safety cabinet using sterile technique, hind legs were disarticulated and placed in RPMI with 10% Fetal Bovine Serum (FBS), 1% penicillin/streptomycin (P/S). Bones were cleaned and placed briefly into 100% ethanol. Bones were then cut on both ends and flushed with RPMI with 10% FBS, 1% P/S using a 27-gauge needle. Marrow was disaggregated and then passed through a 100 µm filter to remove debris. Cells were counted on a hemocytometer using Trypan blue. BM from two hind legs was resuspended in 1 mL PBS supplemented with 1 mM EDTA, 2 % FBS. Monocytes were then isolated using the EasySep Monocyte Isolation Kit (STEMCELL) as per manufacturer recommendations. This method consistently led to isolation of approximately 1-2 x 10^6^ cells with 90-95% Ly6C^hi^ monocytes with no difference in purity between conditions.

### Monocyte adoptive transfer

Monocytes from post-CLP, unoperated, and *Ccr2^-/-^* control mice were isolated and pooled (n = 2-5 mice). *Ccr2^-/-^* recipient mice were anesthetized with ketamine and xylazine and 1×10^6^ monocytes were administered in 100-200 μl of PBS via tail vein injection. Mice then immediately received i.n. LPS to induce lung injury. Treatments were randomized within cage.

### Monocyte and neutrophil ex vivo culture

Monocytes were isolated from BM and immunomagnetically enriched as above. Monocytes from individual post-CLP and unoperated control mice were resuspended at 2.5 x 10^5^ cells/mL in 2 mL DMEM with 10% FBS with LPS (1 μg/mL) in a 12-well plate and incubated for 5 hours at 37 □C with 5% CO_2_. Plates were removed and placed on ice, supernatants removed and centrifuged at 450 g for 10 minutes at 4 □C. Cell-free supernatants were stored at -20 □C until use.

Neutrophils were isolated from BM as previously described^86^. Briefly, whole BM was isolated from hind legs, red cells were lysed using ACK lysis buffer for 1 minute in 37 □C, and cells were washed with RPMI with 10% FBS, 1% P/S. A density gradient was created layering 3 mL of each of the following: Histopaque 1119 (Sigma-Aldrich, density 1.119 g/mL), Histopaque 1077 (Sigma-Aldrich, density 1.077 g/mL), and whole BM from two hind legs in ice cold PBS. Centrifugation was performed at 872 g at room temperature without brake for 30 minutes.

Neutrophils were collected from the Histopaque interface and washed twice with RMPI with 10% FBS with 1% penicillin/streptomycin. Neutrophils from 6 naïve mice were pooled. Cells were counted by trypan blue exclusion on a hemacytometer and purity was assessed to be >90% using DiffQuick (Baxter) on cytospins. Neutrophils were resuspended at 5 x 10^6^ cells/mL in DMEM with 10% FBS and transferred to wells containing 1 mL of thawed monocyte conditioned media (post-CLP or control) or fresh basal media with or without LPS (1 μg/mL) in a 12-well plate. Cells were incubated for 5 hours at 37 □C with 5% CO_2_. Supernatants were isolated as above.

### Serum collection and complete blood cell counts

Whole blood was obtained by puncture of the right ventricle using a heparinized 1 mL syringe and a 26-gauge needle. Needles were removed, blood ejected, and then placed on ice for no more than 1 hour. Samples were centrifuged at 2000g for 10 minutes, with serum stored at -80 ^°^C until time of use. For complete blood cell counts, un-heparinized whole blood was placed into EDTA Microtainer tubes (BD). Blood counts were performed on a Hemavet HV950 (Drew Scientific) in the In-Vivo Animal Core at the University of Michigan.

### Bronchoalveolar lavage, cell count, and differential

Mice were euthanized with ketamine and xylazine. Bronchoalveolar lavage (BAL) was performed as described previously^12^. The trachea was exposed and intubated using a 1.7-mm outer diameter polyethylene catheter. Active insufflation of PBS with 5 mM EDTA in 1 ml aliquots was performed, 3 times per mouse. BAL was centrifuged, supernatant removed, aliquoted, and stored at -80 ^°^C until further use. Cell pellets were resuspended and counted using Trypan blue exclusion counting on a hemocytometer. Cytospins were prepared and stained with Diff-Quick (Baxter) to determine differential for polymorphonuclear nuclear (PMN) and mononuclear cells.

### Flow cytometry and cell sorting

BM or BAL cells were washed, blocked for non-specific staining with anti-CD16/32 Fc receptor block (clone 2.4G2, BD), and stained with fluorophore conjugated antibodies prior to analysis on a Attune NxT (ThermoFisher) or MA900 (Sony) flow cytometer and cell sorter. For BM preparations, cells from a single femur and tibia were lineage depleted using streptavidin beads (STEMCell) and biotinylated antibodies against TER-119, CD19, CD3e. Briefly, cells were resuspended in PBS with 1 mM EDTA, 2% FBS, blocked with 5% rat serum, and incubated with biotinylated antibodies for 5 minutes at room temperature. Lineage depletion was performed on an EasyEights EasySep magnet (STEMCell) per protocol. In some studies evaluating BM progenitors, CD16/32 block was substituted with fluorophore conjugated CD16/32 antibody.

Anti-bodies included: Ly6G (clone 1A8, Biolegend), CD11b (clone M1/70, Biolegend), CD45 (clone 30-F11, Biolegend), Ly6C (clone HK1.4, Biolegend), CD11c (clone N418, Biolegend), CD64 (clone X54-5/7.1.1, BD), CD24 (clone M1/69, BD), MHCII/I-A/E (clone M5/144.15.2, Biolegend), Siglec-F (E50-2440, BD Pharmingen), CD34 (HM34, Biolegend) CD115 (AFS98, Biolegend), CD135/Flt3 (A2-F10, Biolegend), CD117/c-kit (2B8, Biolegend), CD16/32 (93, Biolegend). Lineage markers TER-119 (TER-119, Biolegend), CD3e (145-2C11, Biolegend), B220 (RA3-6B2, Biolegend), CD19 (eBIO1D3, eBioscience) NK1.1 (PK136, Biolegend). 7-AAD was used for live/dead discrimination. Selected populations were isolated by cell sorting, 13,000 - 300,000 cells directly into TRIzol LS (Invitrogen) for RNA-sequencing (RNAseq) and 50 - 60,000 cells into PBS with 2% BSA (Thermofisher) for ATAC-sequencing (ATACseq).

### Quantification of BAL and supernatant proteins

BAL samples and supernatants were assayed for various proteins using the following ELISAs at dilutions where all samples were within the dynamic range of the standard curve: TNFα, IL-1β, IL-6, MIP-2, MCP-1, KC, IL-1ra, S100A8/A9, RAGE (Mouse Duoset, R&D Systems); Pierce BCA assay (Thermo Scientific); Albumin (Bethyl laboratories).

### RNA Isolation

RNA was isolated from monocyte and progenitor populations using TRIzol LS (Invitrogen) as described by the manufacturer. DNAse treatment was performed on RNeasy Mini Kit columns (Qiagen) prior to sequencing.

### RNA-sequencing and analysis

RNA quality was assessed using a bioanalyzer (Agilent) and all RIN were greater than 7. Library preparation, sequencing, and identification of transcript frequency were performed by the University of Michigan Advanced Genomics Core. Library preparation was performed using the low input SMARTer smRNA-Seq kit (Takara). Samples were subjected to 151bp paired-end sequencing, with an average of 30-40 million reads per sample, according to the manufacturer’s protocol (Illumina NovaSeq). BCL Convert Conversion Software (v3.9.3, Illumina) was used to generate de-multiplexed Fastq files. The reads were trimmed using *Cutadapt* (v2.3)^87^. The reads were evaluated with *FastQC* (v0.11.8, Babraham Bioinformatics) to determine quality of the data. Reads were mapped to reference genome GRCm38 (ENSEMBL 102), using *STAR* (v2.7.8a)^88^ or *HISAT2* (v 2.1.0)^89^. Count estimates performed with *RSEM* (v1.3.3)^90^ or *htseq-count* (version 0.13.5)^91^. Alignment options followed ENCODE standards for RNA-seq. PCA was performed using *prcomp* and a single outlier in the CLP group was excluded from analysis for a total of 4 CLP and 5 unoperated samples (Figure S10). Differential gene expression analysis was conducted with *DESeq2* (version 1.38.3)^92^, followed by pathway analysis using *pathfindR* (v1.6.4)^93^.

### ATAC-sequencing and analysis

Cell suspensions were brought to the University of Michigan Advanced Genomics Core for library preparation using the Omni-ATAC-Seq protocol^94^. Libraries were cleaned using MinElute (Qiagen) columns and AMPure XP beads (Beckman Coulter) before quantitation with the Qubit HS dsDNA kit (ThermoFisher), and quality assessment on a TapeStation HS D1000 kit (Agilent). The libraries were pooled and quantitated by qPCR using a Library Quantitation Kit – Illumina Platforms (KAPA) before sequencing on a NextSeq2000 P3 flow cell with an average of 100 million reads per sample. Sample quality was assessed by *FastQC* (v0.11.8, Babraham Bioinformatics). We aligned reads to mm10 with *HISAT2* (v2.1.0)^89^. Filtering steps are performed with *samtools* (v1.2)^95^.Unmapped reads and alignments below a MAPQ threshold were removed. Reads in blacklisted regions (ENCODE Blacklist Regions) were removed using *bedtools* (v2.28.0)^96^. *F-Seq2* was used to call sample-wise peaks (v2.0.3)^97^. Peaks over all samples are merged with *bedtools,* keeping peaks that occurred in at least 3 samples within a treatment group.. Mitochondrial reads were filtered out. Differential peak analysis and pathway enrichment performed using the polyenrich function in *chipenrich* (v2.22.0)^98^. Promoter- and enhancer-specific peaks were determined in R from polyenrich output, <|1000| dist_to_tss and >|1000| dist_to_tss, respectively. Motif enrichment analysis performed using Simple Enrichment Analysis package (*SEA*; v5.5.4)^99^. Finding Individual Motif Occurrences (*FIMO*; v5.5.4) was used to assess AP-1 specific motif enrichment^100^.

### Generation of monocyte subset gene signatures for monocytes and their progenitors

Gene sets for MDP- and GMP-derived monocyte-committed progenitors (MP and cMoP, respectively) and Ly6C^hi^ monocyte subsets (neutrophil-like and DC-like, respectively) were defined by a prior ex vivo experiment^33^. To calculate the enrichment of monocyte and progenitor subset gene signature expression in sorted Ly6C^hi^ monocyte (Lin^-^/Ly6G^-^/CD115^+^/CD11b^hi^/Ly6C^hi^/c-Kit^-^/Flt3^-^) and BM progenitor, gene expression data were normalized with *DESeq2* (v1.38.3) and the mean expression level of all genes in each signature was calculated for each sample and log2 transformed. A Wilcoxon test was used to determine the difference between groups using *ggpubr* (v 0.6.0).

### Human monocyte analysis

Human-mouse conserved monocyte subsets were defined using scRNAseq gene lists from mouse and human lung tumor^68^ (See Genomic Supplement for detailed gene lists). Similarly, mouse-specific signatures were defined using mouse BM Ly6C^hi^ monocytes^36^. Gene transcripts per million (TPM) were used from CD14+ monocytes from critically-ill patients with sepsis^65^ (GSE139913), and hospitalized patients with community acquired pneumonia^66^ (GSE160329), and their respective healthy controls were acquired using the *Gene Expression Omnibus* web client, GEO2R (NCBI, https://www.ncbi.nlm.nih.gov/geo/geo2r/ accessed January 25^th^, 2024). Human peripheral blood monocyte subsets (MS1-MS4) during acute bacterial infection were previously determined^64^. This data was accessed and analyzed through the Single Cell Portal (Broad Institute, https://singlecell.broadinstitute.org/single_cell/study/SCP548/, accessed January 25^th^, 2024) using the gene search and dot plot functions at default settings.

Proteomics data from CD14^+^ monocytes from patients with septic shock were downloaded from the PRIDE repository (PXD023938)^67^. The dataset was acquired in data-dependent acquisition (DDA) mode. MS raw files were converted to mzML format using MSConvert from the ProteinWizard suite. The mzML files were analyzed with the FragPipe computational platform (v23) using the default LFQ_MBR workflow. MSFragger^101^ (v.4.2) was used to search the data against the mouse reference proteome database from Uniprot (downloaded on 2025-04-22), appended with an equal number of reserved sequences and common contaminants. The search results were further processed using MSBooster^102^ for deep learning-based rescoring, Percolator^103^ for PSM validation, ProteinProphet^104^ for protein inference, and Philosopher^105^ for false discovery rate (FDR) filtering at 1% FDR for PSM, peptide and protein levels. The resulting files were passed to IonQuant^106^ to extract and quantify peptides and proteins from the DDA data. The combined protein output file was used for downstream analysis. Only proteins with quantification values in 2 of 3 replicates per sample and present in at least 2 samples per condition were included for the final analysis. Certain proteins (e.g., IL-1R2) were only detected in healthy or sepsis patients and imputation was performed for missing values using the lowest log_2_ protein abundance within the dataset. Enrichment for monocyte subset proteins was tested using MANOVA or PERMANOVA (if imputation performed) using the *stats* and *vegan* R packages between conditions.

### Blood count analysis in patients surviving sepsis

The U.S. Veterans Affairs (VA) healthcare system provides comprehensive medical care to over 6 million veterans^107^. During the study period, VA used a single electronic health record (EHR) system, archived in the Corporate Data Warehouse (CDW) accessible for research^107^. Patient and hospitalization data, including demographics, comorbidities and hospital treatments were extracted from the VA CDW, as described previously^63,108^. Sepsis hospitalizations with live discharge and relevant laboratory data were identified across 138 nationwide VA hospitals (2013 to 2018). Sepsis hospitalizations were identified using electronic health record data, as previously described^9,63,108–110^. Specifically, we identified hospitalizations admitted through the 1) emergency department with evidence of 2) suspected infection and 3) ≥ 1 acute organ dysfunction^109^ with specific criteria available in Supplemental Table 2. Live discharge was defined as being alive on the calendar day of discharge and the day following (to exclude patients discharged home at end of life). We defined relevant laboratory data as having both absolute monocyte count (AMC) and absolute neutrophil count (ANC) on the calendar day of discharge or day prior. Laboratory values were cleaned and standardized as previously described^9,63^. Non-physiologic laboratory values were excluded (Supplemental Table 3) and then the top and bottom 1% of the remaining values were excluded (Supplemental Table 4).

The association of each parameter with 90-day mortality was evaluated through fitting Cox proportional hazards models and restricted cubic splines using the R package *rms* (v6.0-0). To evaluate the effect of concomitant ANC on the association of AMC with 90-day mortality, the interaction between ANC and AMC was modeled using Cox proportional hazards and the interaction surface was plotted using the R package *visreq* (v2.7.0). All statistical code is available online (https://github.com/CCMRPulmCritCare/MonocyteAnalysis).

### Sex as a biological variable

The patient cohort study included female patients. The animal studies were performed in male mice to limit sample size due to inter-animal variability in lung injury response because of prior sepsis. Sex as a biological variable was not evaluated in this study. We have found that female post-CLP mice have similarly enhanced lung injury and prior studies have not shown sex-specific functions of monocytes in LPS-induced lung injury models, to our knowledge, as such we expect the mechanisms to be broadly relevant to both sexes.

### Statistics

Analyses included ANOVA followed by post-hoc testing when ANOVA was significant or unpaired t testing as indicated in the text. In order to minimize spurious comparisons, we prespecified post-hoc comparisons only between CLP/unoperated. Heatmaps were created using standardized Z scores calculated by gene or protein across conditions. All figures show mean and standard error unless otherwise specified. Statistical analyses were carried out in Prism (version 9, Graphpad).

### Study Approval

The patient cohort study was reviewed by the Veterans Affairs Ann Arbor institutional review boards and was deemed exempt from the need for consent under 45 CFR §46, category 4 (secondary use of identifiable data). The murine study protocol was approved by the University Committee on the Use and Care of Animals of the University of Michigan (protocol number 00008999 and 00010712).

## Data Availability

Data is available through the publicly accessible Gene Expression Omnibus (GSE268367 and GSE268368), please contact the corresponding author for other data available upon request. All code is available online at https://github.com/B-McBean/denstaedt_monocyte.

## Author Contributions

SJD, APB, HCP, BBM, HSG, RLZ designed the study. SJD, JC, BA, MWN collected the data. SJD, APB, BM, HCP, HSG, BHS, RLZ analyzed and interpreted the data. SJD, HSG, and RLZ drafted the manuscript. All authors provided critical revision and approval of the final version of this manuscript.

## Acknowledgement

SJD would like to acknowledge Dr. Theodore J. Standiford for his support the concepts of this project. Linda G. Denstaedt for her support. This study was supported (in part) by research funding from National Institutes of Health grants K08HL153799 (to SJD), U01HG011952 (to APB), T32HG000040 (to BM), R35HL144481 (to BBM), R01AG074968 (to BHS), AHRQ R01HS026725 and VA HSR 20-313 (to HCP), R01AI134987 (to HSG), R01HL147920, R01HL131608, R35HL160770 (to RLZ).

