## Supplemental Data for "Long-term immune reprogramming of classical monocytes with altered ontogeny mediates enhanced lung injury in sepsis survivors"

**Supplemental Information**

**Alterations in Monocyte Reprogramming and Ontogeny Enhance Lipopolysaccharide-Induced Lung Injury in Sepsis Survivor Mice**

**Scott J. Denstaedt, Breanna McBean, Alan P. Boyle, Brett C. Arenberg, Matthias Mack, Bethany B. Moore, Michael W. Newstead, Yamei Deng, Alexey I. Nesvizhskii, Benjamin H. Singer, Jennifer Cano, Hallie C. Prescott, Helen S. Goodridge, Rachel L. Zemans**

Table of Contents

| Supplemental Data |  | Page |
| --- | --- | --- |
| Figure S1 | Exudate monocyte/macrophage flow cytometry panel | 2 |
| Figure S2 | Exudate monocyte/macrophage depletion efficacy | 3 |
| Figure S3 | BAL inflammation after monocyte depletion | 4 |
| Figure S4 | BAL inflammation after monocyte adoptive transfer | 5 |
| Figure S5 | Bone marrow monocyte and progenitor flow sorting panel | 6 |
| Figure S6 | Control monocyte ATACseq – pathway enrichment | 7 |
| Figure S7 | Myeloid progenitor flow cytometry gating strategy | 8 |
| Figure S8 | Consort flow diagram for VA sepsis hospitalizations | 9 |
| Figure S9 | Conserved human-mouse monocyte subset signature | 10 |
| Figure S10 | Principle Component Analysis Ly6C^hi^ monocyte RNAseq | 11 |
| Table S1 | Characteristics of Sepsis Cohort | 12 |
| Table S2 | Sepsis hospitalization identification criteria | 13 |
| Table S3 | Patient laboratory values after trimming non-physiologic data | 14 |
| Table S4 | Patient laboratory values after trimming top and bottom 1% | 14 |
| References | References | 15 |

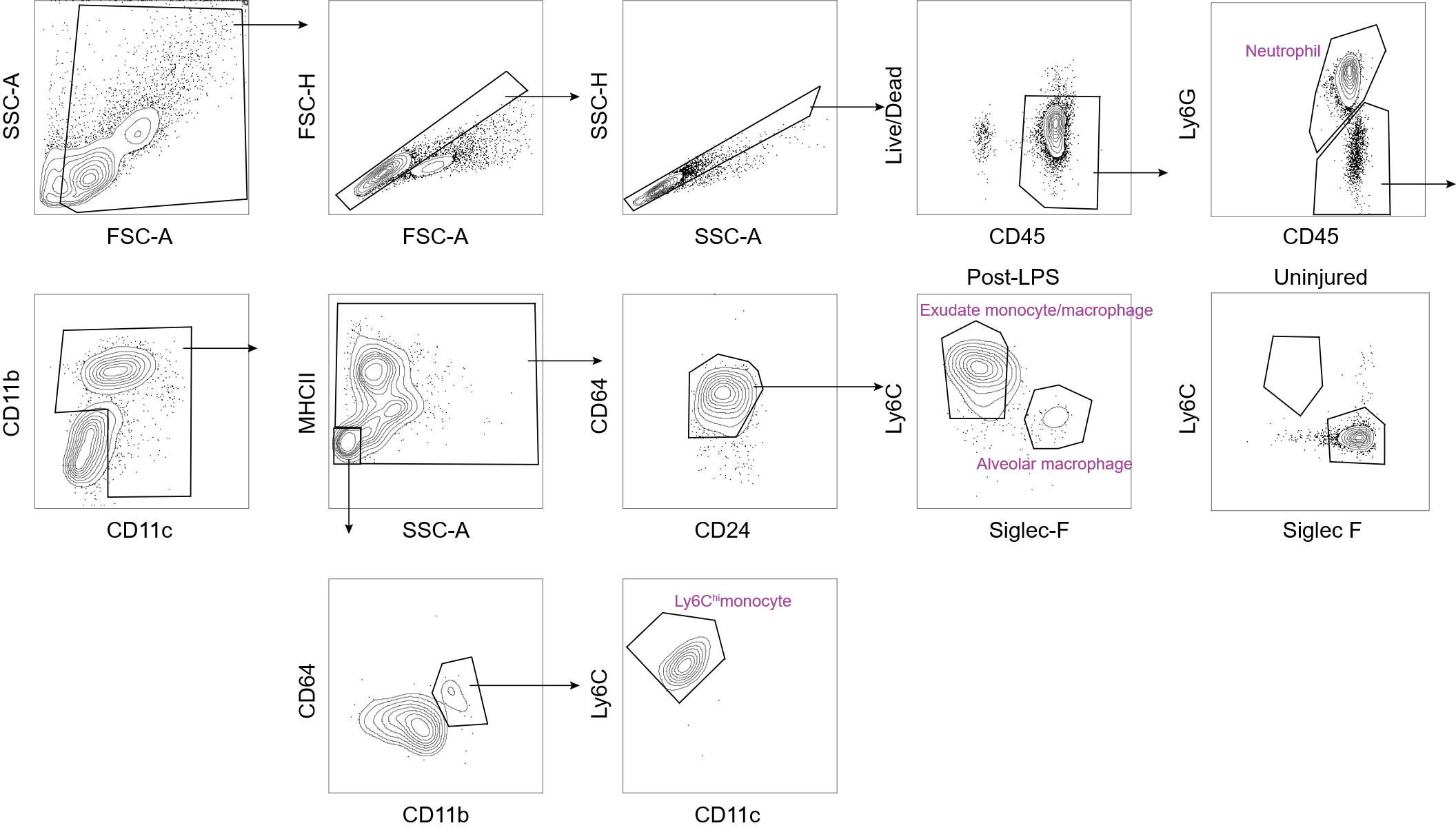

**Figure S1. Exudate monocyte and macrophage flow cytometry panel performed on BAL.** Exudate monocyte/macrophages in the BAL were assessed as previously described ^1^. Exudate monocyte/macrophage were identified within the myeloid (CD11b+ or CD11c+) population as MHCII^+^/SSC^hi^/CD64^+^/Ly6C^+^/Siglec-F^-^ events, and Ly6C^hi^ monocytes were identified within the myeloid population (CD11b^+^ or CD11c^+^) as MHCII^-^/SSC^lo^/CD64/Ly6C^hi^ events. All populations gated using FMO controls. As shown, exudate monocyte/macrophage are not present in the BAL in uninjured lung but are present post-LPS (72 hours) indicating recruitment to the airspace under LPS administration.

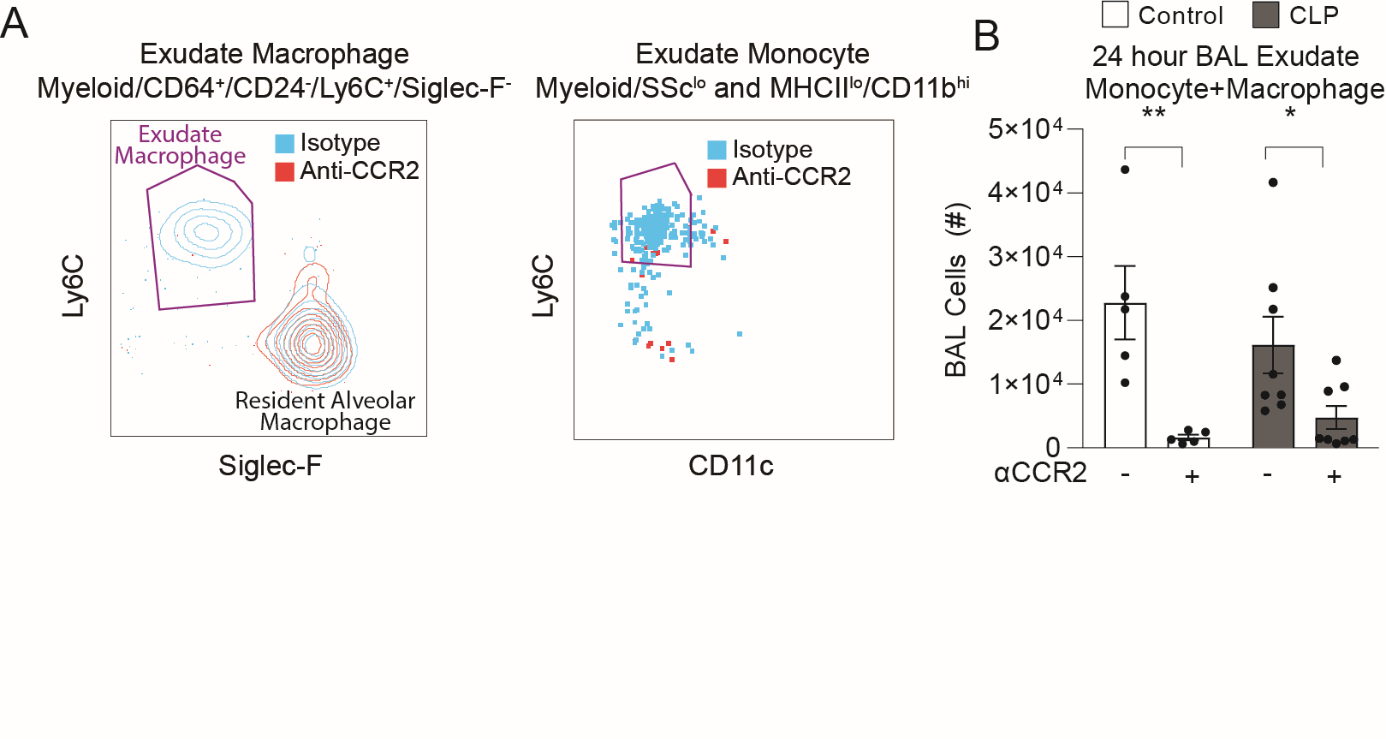

**Figure S2. Anti-CCR2 treatment effectively depletes exudate Ly6C^hi^ monocytes and macrophages.** Exudate monocyte/macrophages were assessed 24 hours after LPS administration following monocyte depletion in BAL fluid (A). Exudate monocyte/macrophage were identified (Figure S1), proportions shown for exudate monocyte/macrophage combined before and after depletion. 1 cohort shown, n = 5-8 per group. Mean ± SEM, post-hoc sidak t-test shown. * p < 0.05, ** p < 0.01. CLP, Cecal Ligation and Puncture. LPS, Lipopolysaccharide. BAL, bronchoalveolar lavage.

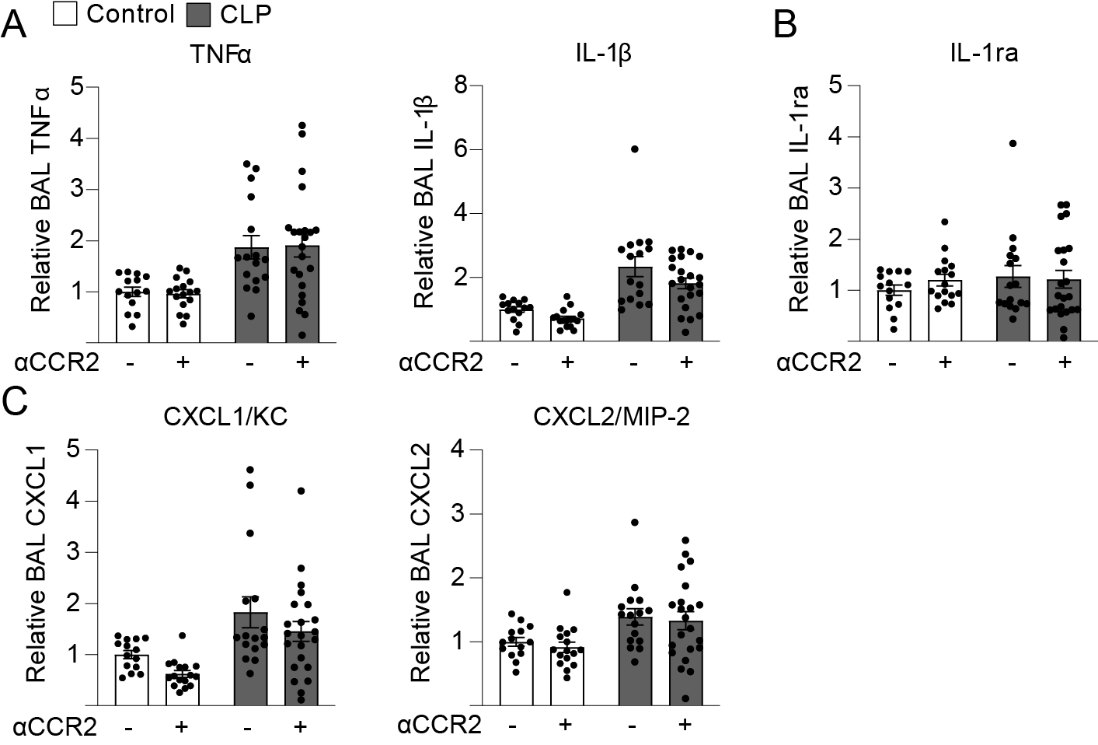

**Figure S3. BAL inflammation following monocyte depletion.** Lung inflammatory proteins were assessed in BAL 72 hours after LPS administration. Inflammatory proteins (A), anti-inflammatory proteins (B), and neutrophil chemokines shown (C). n = 4-8 per group, 3 cohorts. Relative BAL protein levels expressed relative to isotype treated control mice. Mean ± SEM shown. BAL, bronchoalveolar lavage. CLP, cecal ligation and puncture.

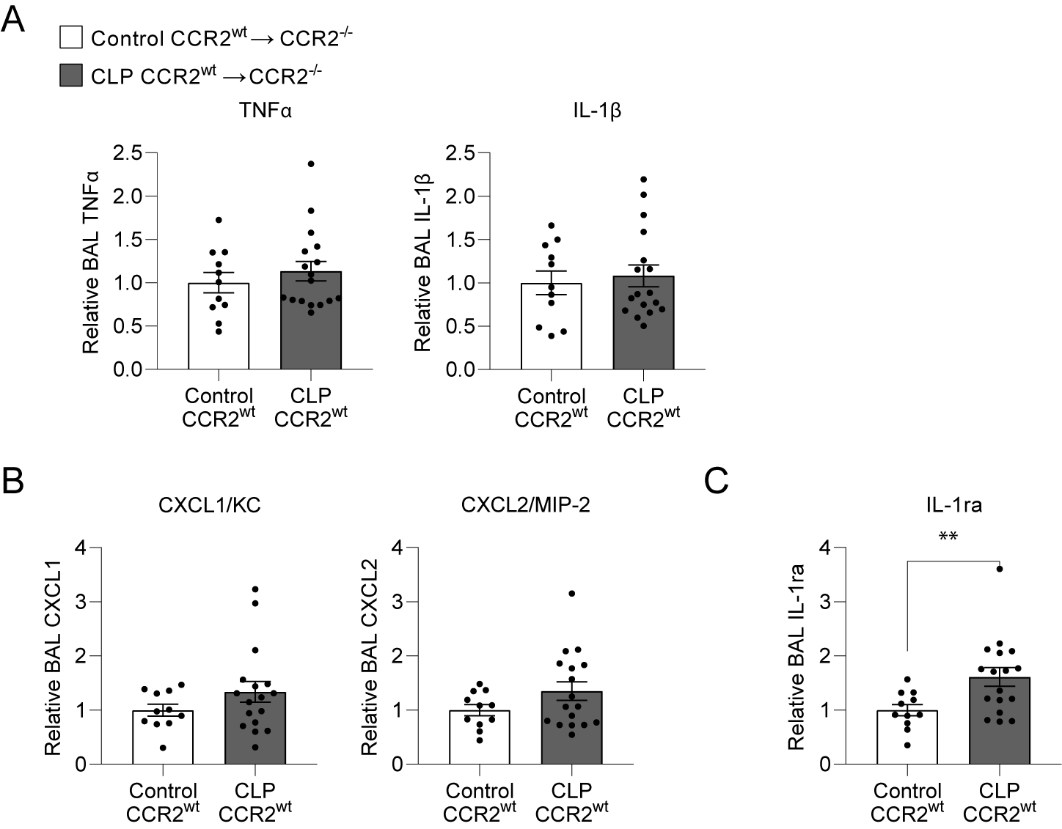

**Figure S4. BAL inflammation following monocyte adoptive transfer.** Lung inflammatory proteins were assessed 72 hours after adoptive transfer of monocytes. Inflammatory proteins (A), neutrophil chemokines (B), and anti-inflammatory proteins shown (C). Protein measurements expressed relative to the CCR2^wt^ unoperated control transfer within each cohort. 3 cohorts shown, n = 3-6 per group. Mean ± SEM, Welch's t-test p-value shown. ** p < 0.01. CLP, Cecal Ligation and Puncture. BAL, bronchoalveolar lavage.

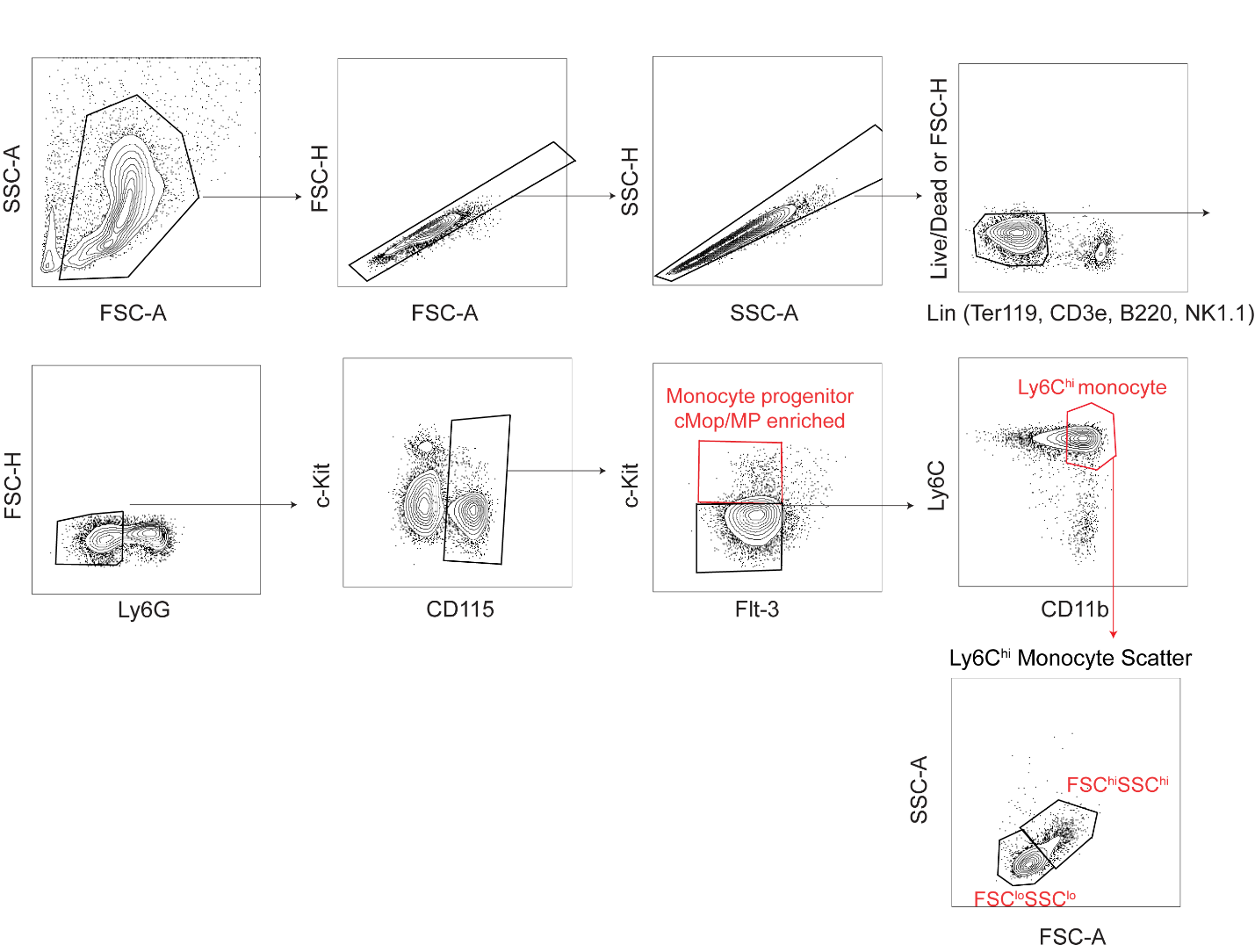
**Figure S5. Bone marrow monocyte and progenitor flow gating strategy for flow sorting.** Ly6C^hi^ monocytes and cMoP/MP populations (red boxes) were identified and sorted as previously described ^2^. All populations gated using FMO controls.

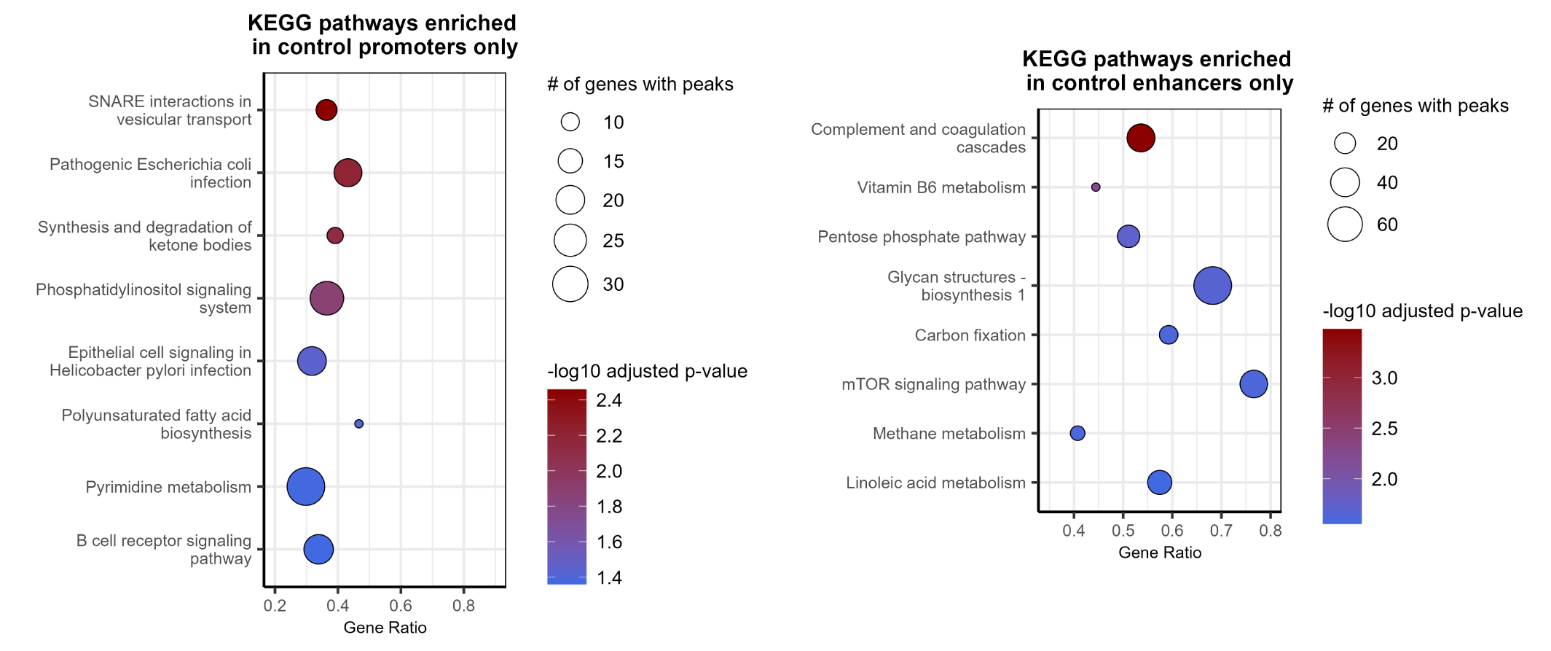

**Figure S6. Chromatin accessibility pathway enrichment in control Ly6C^hi^ monocytes.** Promoters were defined as genomic regions < 1 kb from a transcription start site (TSS). Enhancers were defined as genomic regions > 1 kb from a TSS.

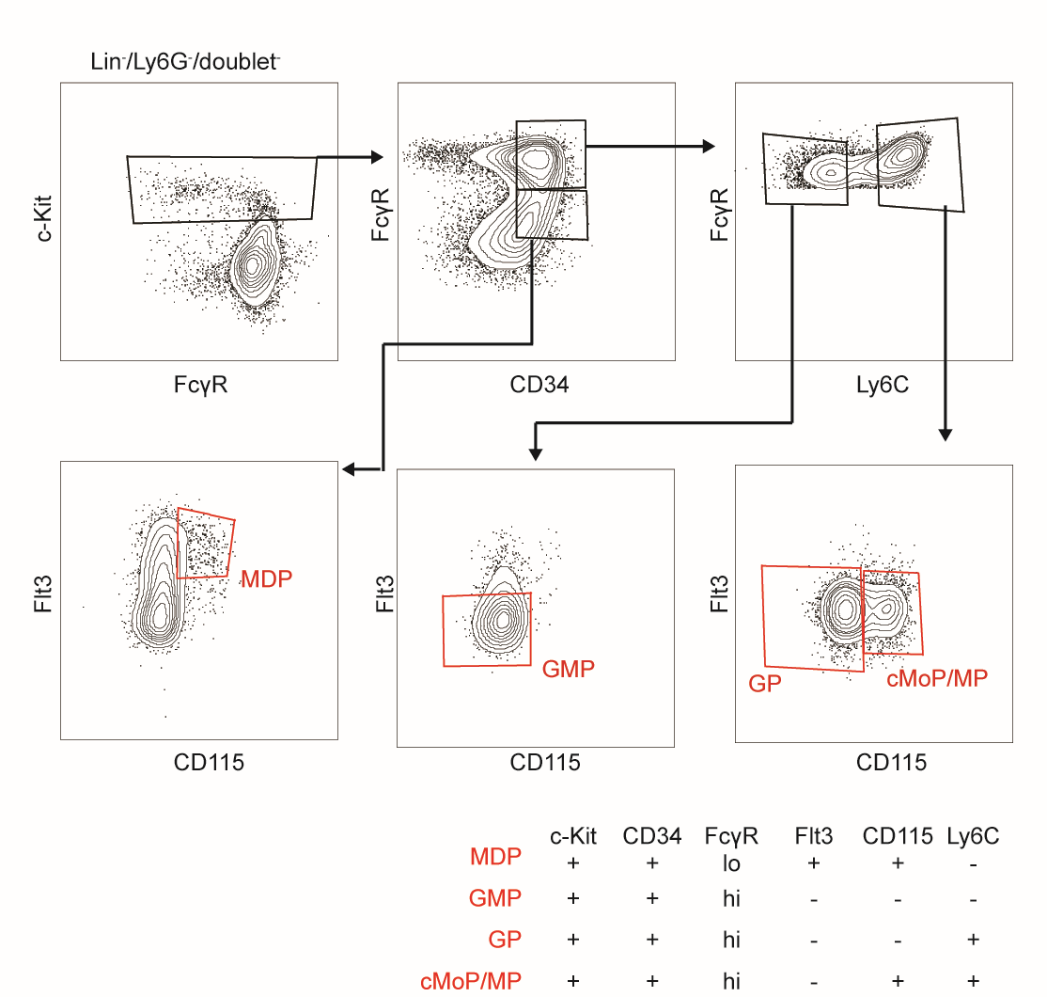

**Figure S7**. **Myeloid bone marrow progenitor flow cytometry gating strategy.** Myeloid progenitors were identified as previously described^3^. All populations gated using FMO controls.

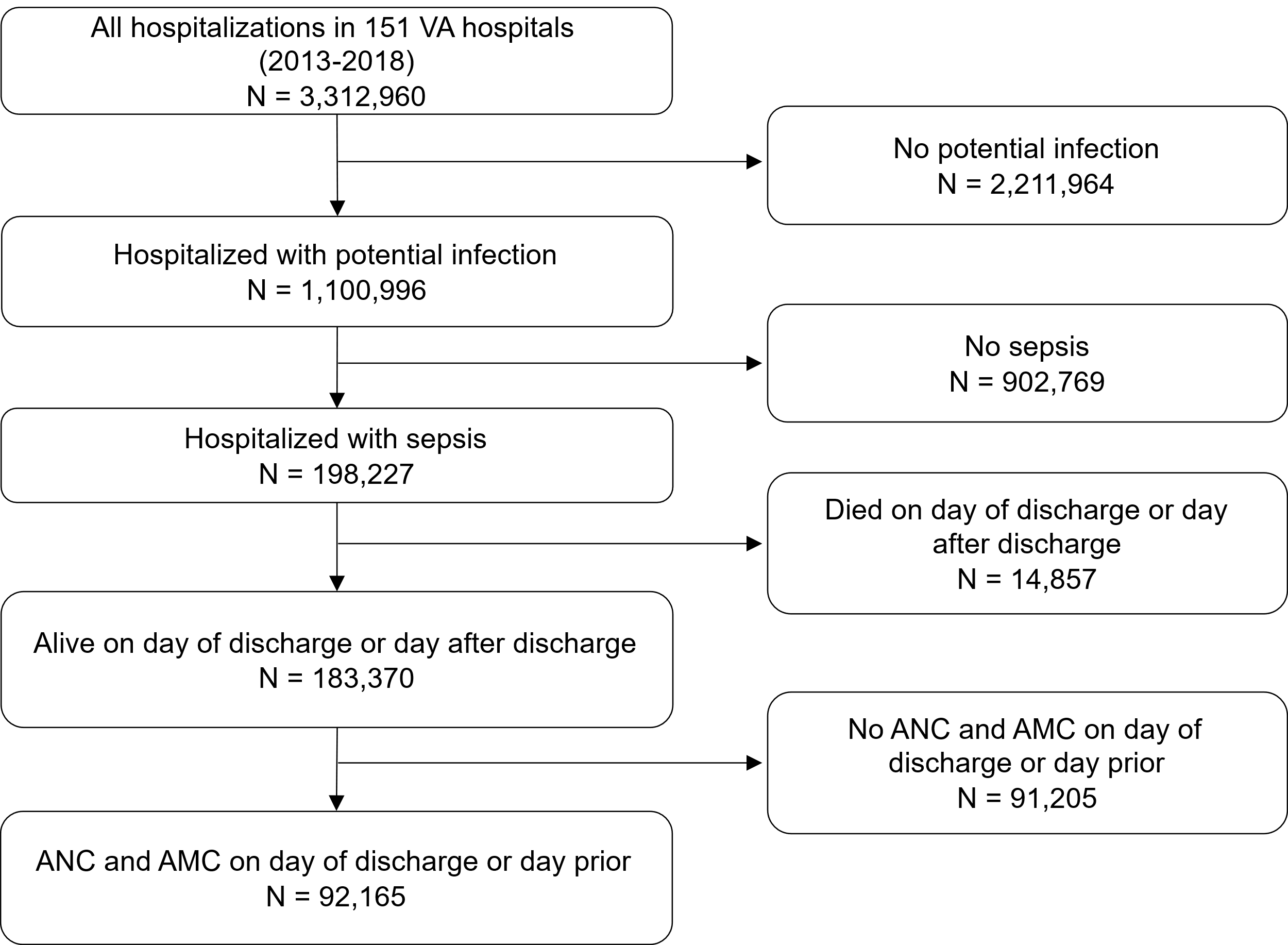

**Figure S8. Consort Diagram of VA Hospitalizations**. Hospitalizations with potential infection were defined as ≥2 SIRS criteria and treatment with antimicrobials within 48 hours of arrival. Hospitalizations for sepsis were defined, as in supplemental Table 1, as potential infection with continuation of antimicrobials for at least 4 days and ≥ 1 acute organ dysfunctions within 48 hours of arrival through the emergency department. ANC, absolute neutrophil count. AMC, absolute monocyte count.

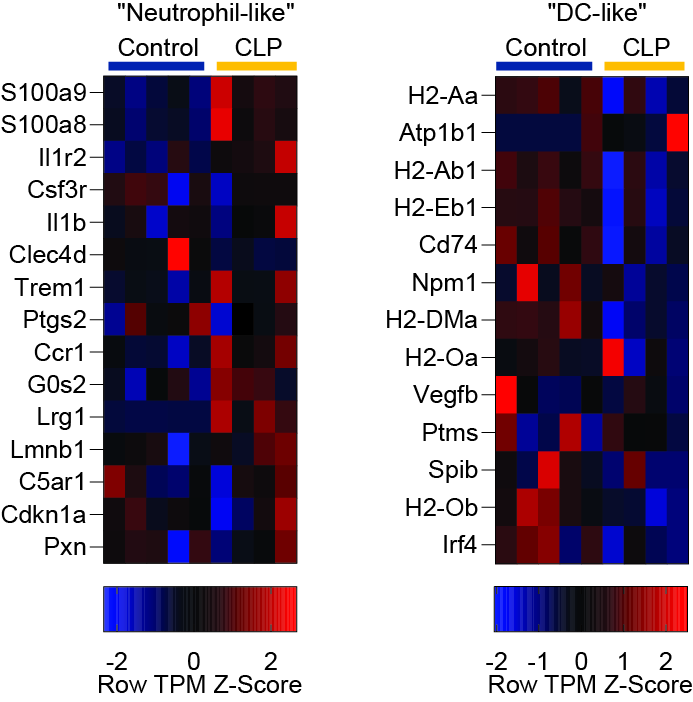

**Figure S9. Conserved human-mouse signature**^4^ **in mouse Ly6C^hi^ monocytes.** Row TPM Z-score shown.

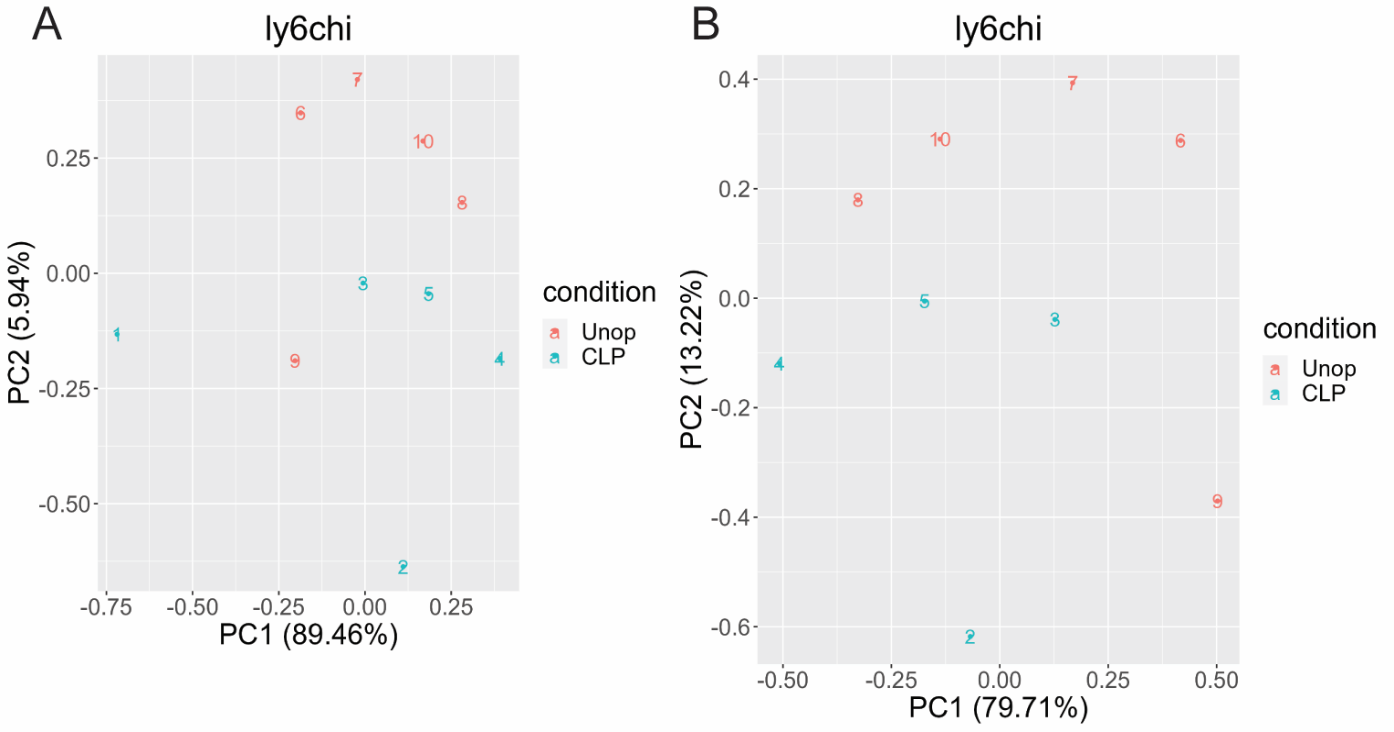

**Figure S10. PCA plot of RNA-seq analysis performed on flow sorted Ly6C^hi^ monocytes.** PCA plot shown before (A) and after (B) exclusion of a single outlier sample in the CLP group (sample #1).

| **Supplemental Table 1. Characteristics of Sepsis Cohort** | | | |
| --- | --- | --- | --- |
|  | **Total Cohort**  n = 92165 (100.0) | **No mortality**  n = 79600 (86.4) | **90-day**  **all-cause mortality**  n = 12565 (13.6) |
| **Age, median (IQR)** | 68 (62, 76) | 68 (61, 75) | 72 (66, 83) |
| **Sex, M, %** | 88811 (96.4) | 76533 (96.1) | 12278 (97.7) |
| **Race, %** |  |  |  |
| Black | 18622 (20.2) | 16221 (20.4) | 2401 (19.1) |
| White | 65932 (71.5) | 56887 (71.5) | 9045 (72.0) |
| Other | 2003 (2.2) | 1782 (2.2) | 221 (1.8) |
| Unknown | 5463 (5.9) | 4605 (5.8) | 858 (6.8) |
| **Comorbidity count, median (IQR)** | 7 (4, 9) | 6 (4, 9) | 8 (6, 10) |
| **Select comorbid conditions, n (%)** |  |  |  |
| Any Diabetes | 47522 (51.6) | 41336 (51.9) | 6186 (49.2) |
| Chronic pulmonary disease | 43377 (47.1) | 36790 (46.2) | 6587 (52.4) |
| CKD | 33130 (35.9) | 27592 (34.7) | 5538 (44.1) |
| Diabetes with complication | 31843 (34.5) | 27683 (34.8) | 4160 (33.1) |
| Congestive heart failure | 29992 (32.5) | 24636 (30.9) | 5356 (42.6) |
| Any Cancer | 23721 (25.7) | 18338 (23) | 5383 (42.8) |
| Liver disease | 17237 (18.7) | 14315 (18) | 2922 (23.3) |
| Neurologic disease | 16561 (18.0) | 13463 (16.9) | 3098 (24.7) |
| Metastatic cancer | 6646 (7.2) | 4202 (5.3) | 2444 (19.5) |
| Rheumatoid Arthritis/Collagen Vascular Diseases | 4022 (4.4) | 3473 (4.4) | 549 (4.4) |
| **Acute organ dysfunction count, median (IQR)** | 1 (1, 2) | 1 (1, 2) | 1 (1, 2) |
| **Acute organ dysfunctions, n (%)** |  |  |  |
| Renal | 57142 (62.0) | 48894 (61.4) | 8248 (65.6) |
| Elevated lactate | 39523 (42.9) | 33993 (42.7) | 5530 (44.0) |
| Hematologic | 10646 (11.6) | 8544 (10.7) | 2102 (16.7) |
| Hepatic | 11878 (12.9) | 9976 (12.5) | 1902 (15.1) |
| Circulatory (vasopressor use) | 6625 (7.2) | 5377 (6.8) | 1248 (9.9) |
| Respiratory (mechanical  ventilation) | 4093 (4.4) | 3326 (4.2) | 767 (6.1) |
| **ICU admission, n (%)** | 26513 (28.8) | 21745 (27.3) | 4768 (37.9) |
| **Hospital LOS, median (IQR, days)** | 6 (4, 10) | 6 (4, 9) | 8 (5, 12) |
| **Discharge blood parameter, median (IQR)** |  |  |  |
| ANC (10^3^/µL) | 5.64 (3.83, 7.88) | 5.55 (3.80, 7.70) | 6.38 (4.20, 9.04) |
| AMC (10^3^/µL) | 0.70 (0.50, 0.93) | 0.70 (0.50, 0.93) | 0.70 (0.47, 0.97) |
| Comorbidity count determined using 30 Elixhauser comorbidities ^5^.  ANC, absolute neutrophil count. AMC, absolute monocyte count. | | | |

| ­­Supplemental Table 2. Sepsis Hospitalization Criteria – Veterans Affairs Hospital Data | | |
| --- | --- | --- |
| **Broad Criteria^‡^** | **Specific Criteria** | **References** |
| Admission through the emergency department | N/A |  |
| Suspected infection | 2+ systemic inflammatory response criteria (SIRS), | ^6,7^ |
|  | Initiation of systemic antimicrobials (oral or intravenous) within 48 hours of ED arrival, |  |
|  | AND Continuation of antimicrobials for at least 4 days, inclusive of antibiotics prescribed at discharge to be completed at home |  |
| ≥ 1 acute organ dysfunction within 48 hours of emergency department arrival | acute renal dysfunction^†^ | ^6^ |
|  | acute liver dysfunction^†^ |  |
|  | acute hematologic dysfunction^†^ |  |
|  | lactate > 2.0 mmol/L |  |
|  | receipt of invasive mechanical ventilation |  |
|  | receipt of systemic vasopressors |  |
| ^†^Acute renal, liver, and hematologic dysfunction were defined as worsening of creatinine, total bilirubin, or platelet count respectively from baseline. Baseline was defined as the best laboratory value during hospitalization or the preceding 6 months ^6^.  ^‡^Broad criteria for sepsis hospitalization are adapted from the CDC definition for an adult sepsis event (ASE) | | |

| **Supplemental Table 3: Lab Data Check Table (Pre-Trim)**:  Use the last CBC parameters of hospitalization that occur on day of discharge or day prior,  AFTER excluding non-physiologic labs (values ≤ 0 or ≥ 300 x 10^3^ cells/μl)  BEFORE excluding top/bottom percentile | | | | | | | | | | |
| --- | --- | --- | --- | --- | --- | --- | --- | --- | --- | --- |
| **CBC Parameter** | **Total**  **measurements** | **% of hospitalizations** | **Mean** | **Min** | **1^st^%ile** | **25%ile** | **Median** | **75%ile** | **99^th^%ile** | **Max** |
| ANC | 103171 | 56.3 | 6.27 | 0.00 | 0.10 | 3.80 | 5.65 | 7.96 | 18.50 | 195.10 |
| AMC | 104020 | 56.7 | 0.76 | 0.01 | 0.08 | 0.50 | 0.70 | 0.95 | 2.10 | 135.50 |

| **Supplemental Table 4: Lab Data Check Table (POST-Trim)**:  Use the last CBC parameters of hospitalization that occur on day of discharge or day prior,  AFTER excluding non-physiologic labs  AFTER excluding top/bottom percentile | | | | | | | | | | |
| --- | --- | --- | --- | --- | --- | --- | --- | --- | --- | --- |
| **CBC Parameter** | **Total**  **measurements** | **% of hospitalizations** | **Mean** | **Min** | **1^st^%ile** | **25%ile** | **Median** | **75%ile** | **99^th^%ile** | **Max** |
| ANC | 101324 | 55.3 | 6.12 | 0.10 | 0.50 | 3.80 | 5.63 | 7.90 | 15.91 | 18.50 |
| AMC | 102130 | 55.7 | 0.74 | 0.08 | 0.10 | 0.50 | 0.70 | 0.93 | 1.80 | 2.10 |
